# Adaptive protein synthesis in genetic models of copper deficiency and childhood neurodegeneration

**DOI:** 10.1101/2024.09.09.612106

**Authors:** Alicia R. Lane, Noah E. Scher, Shatabdi Bhattacharjee, Stephanie A. Zlatic, Anne M. Roberts, Avanti Gokhale, Kaela S. Singleton, Duc M. Duong, Mike McKenna, William L. Liu, Alina Baiju, Felix G Rivera Moctezuma, Tommy Tran, Atit A. Patel, Lauren B. Clayton, Michael J. Petris, Levi B. Wood, Anupam Patgiri, Alysia D. Vrailas-Mortimer, Daniel N. Cox, Blaine R. Roberts, Erica Werner, Victor Faundez

## Abstract

Rare inherited diseases caused by mutations in the copper transporters *SLC31A1* (CTR1) or *ATP7A* induce copper deficiency in the brain, causing seizures and neurodegeneration in infancy through poorly understood mechanisms. Here, we used multiple model systems to characterize the molecular mechanisms by which neuronal cells respond to copper deficiency. Targeted deletion of CTR1 in neuroblastoma cells produced copper deficiency that was associated with a metabolic shift favoring glycolysis over oxidative phosphorylation. Proteomic and transcriptomic analysis of CTR1 KO cells revealed simultaneous upregulation of mTORC1 and S6K signaling and reduced PERK signaling. Patterns of gene and protein expression and pharmacogenomics show increased activation of the mTORC1-S6K pathway as a pro-survival mechanism, ultimately resulting in increased protein synthesis. Spatial transcriptomic profiling of *Atp7a^flx/Y^ :: Vil1^Cre/+^* mice identified upregulated protein synthesis machinery and mTORC1-S6K pathway genes in copper-deficient Purkinje neurons in the cerebellum. Genetic epistasis experiments in *Drosophila* demonstrated that copper deficiency dendritic phenotypes in class IV neurons are partially rescued by increased S6k expression or 4E-BP1 (Thor) RNAi, while epidermis phenotypes are exacerbated by Akt, S6k, or raptor RNAi. Overall, we demonstrate that increased mTORC1-S6K pathway activation and protein synthesis is an adaptive mechanism by which neuronal cells respond to copper deficiency.

**Significance:**

- Copper deficiency is present in rare conditions such as Menkes disease and CTR1 deficiency and in more common diseases like Alzheimer’s. The mechanisms of resilience and ultimate susceptibility to copper deficiency and associated pathology in the brain remain unknown.
- We demonstrate that in a human cell line, *Drosophila*, and the mouse cerebellum, copper-deficient neuronal cells exhibit increased protein synthesis through mTORC1 activation and decreased PERK (EIF2AK3) activity.
- Upregulation of protein synthesis facilitates resilience of neuronal cells to copper deficiency, including partial restoration of dendritic arborization. Our findings offer a new framework for understanding copper deficiency-related pathology in neurological disorders.

## Introduction

Copper is an essential micronutrient but can be toxic if not regulated appropriately (Zlatic et al., 2015; Lutsenko et al., 2024). Neurodevelopment has stringent temporal and spatial requirements for copper levels and localization within organelles and across tissues, and a failure of mechanisms controlling copper homeostasis is causative of or associated with a variety of neurodevelopmental and neurodegenerative diseases (Waggoner et al., 1999; Opazo et al., 2014). Defects in copper homeostasis are linked to dysfunctional neuronal differentiation, organization, migration, and arborization; axonal outgrowth; synaptogenesis; and neurotransmission in multiple brain regions (Purpura et al., 1976; Hirano et al., 1977; Büchler et al., 2003; Niciu et al., 2006; El Meskini et al., 2007; Scheiber et al., 2014; Zlatic et al., 2015).

As copper is a redox active metal required for oxidative phosphorylation, it is tightly linked to bioenergetics (Garza et al., 2022b). Thus, tissues with high energy demands, like the brain, are particularly susceptible to copper toxicity and deficiency (Scheiber et al., 2014). Neurodevelopment is a particularly vulnerable time due to the high energy consumption and transitions in metabolism by the brain during this period (Bülow et al., 2022) and the increased neuronal demand for copper after differentiation (Hatori et al., 2016). While the fetal brain produces energy by glycolysis, after birth there is an increase in brain consumption of glucose and oxygen that peaks in humans at age 5, nearly doubling the consumption of the adult brain (Goyal et al., 2014; Kuzawa et al., 2014; Steiner, 2020; Oyarzábal et al., 2021). This transition from glycolysis to oxidative phosphorylation is in part cell-autonomous, as it is also observed in differentiating neurons and muscle cells in culture (Vest et al., 2018; Iwata et al., 2023; Casimir et al., 2024; Rajan and Fame, 2024), and is necessary for neurodevelopment (Zheng et al., 2016; Sakai et al., 2023; Casimir et al., 2024; Iwata and Vanderhaeghen, 2024). In fact, aberrant hyperglycolytic metabolism in neurons induces dysfunction and damage in vitro and in vivo (Jimenez-Blasco et al., 2024).

Mutations affecting copper homeostasis provide an opportunity to understand how copper-dependent mechanisms and metabolism interact to drive brain development. For example, mutations affecting the copper transporter ATP7A cause conditions of varying severity and age of onset but which all present with prominent neurological symptoms (OMIM: 309400, 304150, 300489; Kaler, 2011). In its most severe form, Menkes disease, overt neurological symptoms and neurodegeneration appear between 1-3 months of age (Menkes et al., 1962; Tümer and Møller, 2010; Kaler, 2011; Skjørringe et al., 2017). Similarly, most Menkes mouse models exhibit substantial neurodegeneration by postnatal days 10 to 14, typically culminating in death by day 21 (Yajima and Suzuki, 1979; Iwase et al., 1996; Donsante et al., 2011; Kaler, 2011; Lenartowicz et al., 2015; Guthrie et al., 2020; Yuan et al., 2022). Whether this delay in disease appearance is impacted by increasing neurodevelopmental demands for copper and/or neurodevelopmental-sensitive mechanisms conferring resilience to copper depletion has not been considered.

Here, we sought to identify cell-autonomous mechanisms in neurons downstream of copper deficiency. We generated *SLC31A1*-null cells that lack the ability to import copper through the plasma membrane copper transporter CTR1. *SLC31A1* genetic defects (OMIM: 620306) cause symptoms similar to Menkes disease (Batzios et al., 2022; Dame et al., 2022). We discovered that these copper-deficient cells exhibit impaired mitochondrial respiration concurrent with increased glycolysis. Using multiomics approaches, we discovered the modification of two distinct pathways involved in regulation of protein synthesis in CTR1 KO cells: increased mTORC1 signaling pathway activation and reduced activity of EIF2AK3 (PERK, eukaryotic translation initiation factor 2 alpha kinase 3). Accordingly, CTR1-null cells had increased protein synthesis as measured by puromycin incorporation. Similarly, copper-deficient *Atp7a^flx/Y^ :: Vil1^Cre/+^* mice upregulate protein synthesis machinery and mTOR pathway transcripts. Based on our pharmacogenomics in cells and genetic epistasis experiments in *Drosophila*, which demonstrate a partial rescue of dendritic phenotypes by S6k overexpression or Thor RNAi, we conclude that mTOR activation and upregulation of protein synthesis is an adaptive mechanism engaged in response to copper deficiency.

## Results

### Metabolic phenotypes in a cell-autonomous model of copper deficiency

To establish a cell-autonomous model of copper deficiency, we generated *SLC31A1* null (hereafter referred to as CTR1 KO) SH-SY5Y clonal cells by CRISPR genome editing (Extended Data Fig. 1-1). *SLC31A1* encodes the copper importer CTR1, for which protein expression was abolished after CRISPR genome editing (Fig. 1A, Extended Data Fig. 1-1A). CTR1 KO clones were characterized by their increased resistance to copper (Extended Data Fig. 1-1B) and reduced abundance of the copper-dependent Golgi enzyme dopamine-β-hydroxylase (DBH, Fig. 1A). We confirmed that relative to wild type cells, whole CTR1 KO cells were selectively depleted of copper but not zinc as measured by inductively-coupled mass spectrometry (ICP-MS) (Wilschefski and Baxter, 2019; Lane et al., 2022) (Fig. 1B, Extended Data Fig. 1-1D). Mitochondrial-enriched fractions were also selectively depleted of copper but not zinc in CTR1 KO cells (Fig. 1B, Extended Data Fig. 1-1D). These whole cell and mitochondrial copper phenotypes were rescued by a non-toxic dose of elesclomol (Fig. 1B, Extended Data Fig. 1-1C), a small molecule that delivers copper preferentially to the mitochondria (Kirshner et al., 2008; Wu et al., 2011; Blackman et al., 2012; Soma et al., 2018; Garza et al., 2022a). As copper is essential for the assembly and function of the electron transport chain, particularly Complex IV, we chose to examine the levels of subunits of respiratory chain complexes as well as assembly of supercomplexes. The protein abundance of the copper-dependent mitochondrial Complex IV in CTR1 KO cells was 55% of wild-type levels, while there was no decrease in levels of the other respiratory complexes (Fig. 1C). Using blue native gel electrophoresis with digitonin to preserve supercomplexes, we demonstrated Complex I, III, and IV organization into complexes was compromised in CTR1 KO cells, with more pronounced changes in Complex III and IV. We did not detect modifications of Complex II (Fig. 1D).

**Figure 1.**
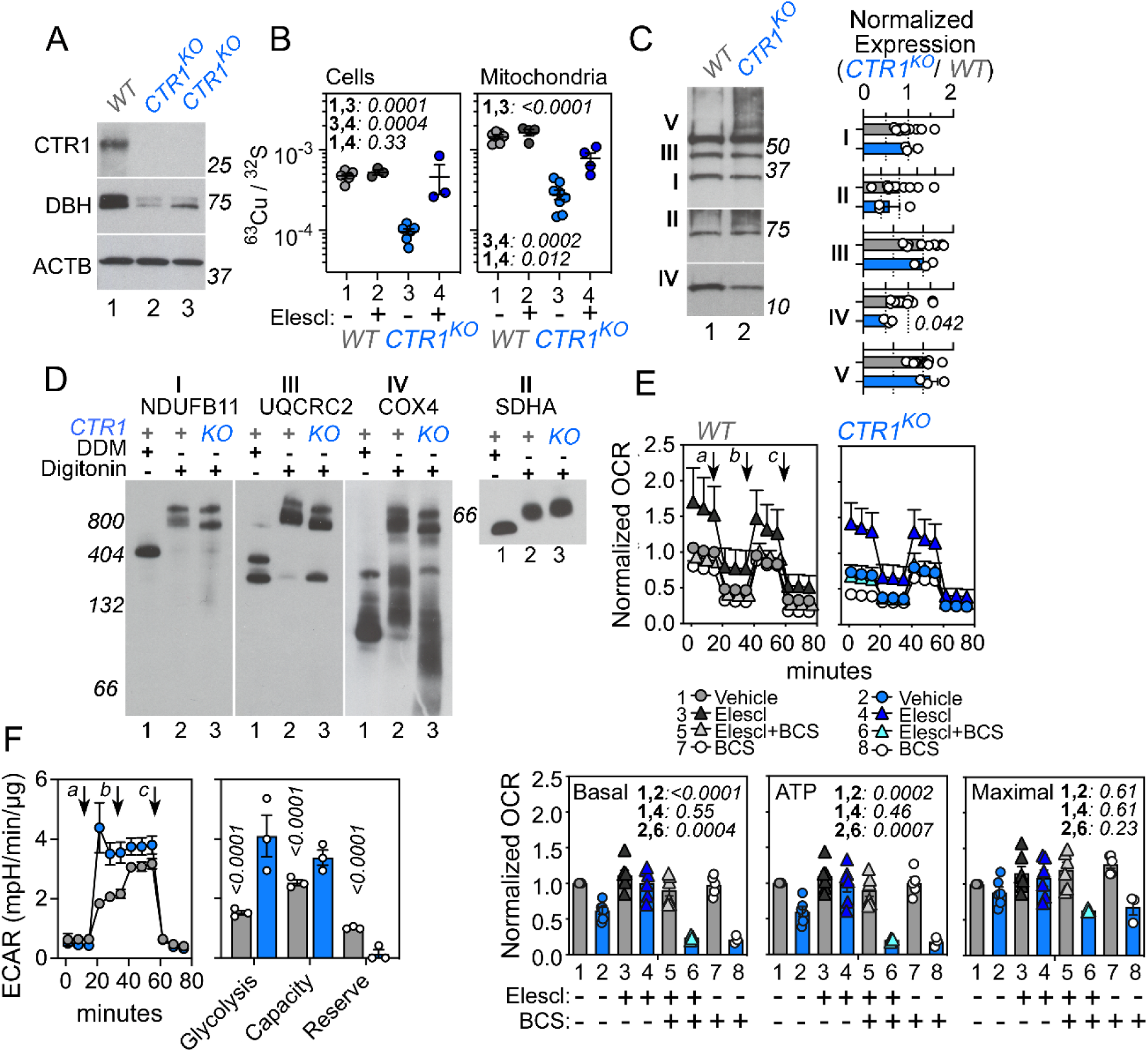
CTR1 (*SLC31A1*) null mutation disrupts electron transport chain assembly and function and increases glycolysis. **A.** Immunoblot of cellular extracts from wild-type (lane 1) and two independent *SLC31A1Δ/Δ* mutant (CTR1 KO, lanes 2-3) SH-SY5Y cell clones probed for CTR1 and DBH with beta-actin as a loading control. **B.** 63Cu quantification in whole cells or mitochondria in CTR1 KO cells treated with vehicle or 1 nM elesclomol, normalized to 32S. Italicized numbers represent q values (One-Way ANOVA, followed by Benjamini, Krieger, and Yekutieli multiple comparisons correction). **C.** Immunoblot with OxPhos antibody mix in mitochondrial fractions from wild-type and CTR1 KO cells. Complex II was used as a loading control as it does not form respiratory supercomplexes (Iverson et al., 2023). (Each dot is an independent biological replicate. Italicized numbers represent p values analyzed by two-sided permutation t-test.) **D.** Blue native electrophoresis of mitochondrial fractions from wild-type and CTR1 KO cells (Clone KO3) solubilized in either DDM or digitonin to preserve or dissolve supercomplexes, respectively (Wittig et al., 2006; Timón-Gómez et al., 2020). Shown are native gel immunoblots probed with antibodies against Complex, I, II, III, and IV. Italicized numbers represent p values. Complex II was used as a loading control. Immunoblots were also prepared with CTR1 clone KO20 (not shown). **E-G.** Seahorse stress tests in wild-type and CTR1 KO cells. Arrows indicate the sequential addition of oligomycin (a), FCCP (b), and rotenone-antimycin (c) in the Mito Stress Test (**E**) to cells treated with vehicle, 1 nM elesclomol, or 200 μM BCS for 72 hours (**E, F**, BCS n =3, all other treatments n=6-7) or the addition of glucose (a), oligomycin (b), and 2-Deoxy-D-glucose (c) in the Glycolysis Stress Test (**G**, n=3). Basal cellular respiration and glycolysis were measured for 90 min after additions using Seahorse. **E**,**F**. Mito Stress Test data are presented normalized to basal respiration of wild-type cells in the absence of drug, analyzed by a One-Way ANOVA followed by Benjamini, Krieger, and Yekutieli multiple comparisons correction (italics show q values). CTR1 clone KO20 was used. **G**. Glycolysis Stress Test data are presented normalized to protein, analyzed by two-sided permutation t-test (italicized numbers represent p values). All data are presented as average ± SEM.

To examine the function of the electron transport chain in CTR1 KO cells, we performed Seahorse oximetry using the Mito Stress Test. Consistent with their reduced levels and impaired assembly of Complex IV, CTR1 KO cells exhibited decreased basal and ATP-dependent respiration (0.53x and 0.54x wild-type levels, respectively; Fig. 1E, F, compare columns 1 and 2). These differences attributable to copper deficiency in CTR1 KO cells since they were magnified by treatment with the cell-impermeant copper chelator BCS (Fig. 1F, compare columns 2 and 8), while BCS had no effect on wild-type cells under these conditions (Fig. 1F, compare columns 1 and 7). Copper delivery via elesclomol treatment at a low, non-toxic concentration rescued the respiration defects and increased media acidification (suggesting increased glycolysis) in CTR1 KO cells back to wild type levels (Fig. 1F, compare columns 1, 2, and 4, Extended Data Fig. 1-1C,E). BCS suppressed the elesclomol-mediated rescue of respiration and acidification phenotypes in CTR1 KO cells (Fig. 1F, compare columns 4 and 6, Extended Data Fig. 1-1E), confirming the requirement for copper in this rescue (Extended Data Fig. 1-1E). There was no difference in maximal respiration in untreated cells or in response to BCS or elesclomol (Fig. 1F). The increased media acidification by CTR1 KO cells suggests increased glycolysis as compared to wild type cells (Extended Data Fig. 1-1E). To characterize the glycolytic parameters of CTR1 KO cells, we used the Glycolysis Stress Test. CTR1 KO cells exhibited elevated extracellular acidification rate, with increased glycolysis (2.03x wild-type levels), increased glycolytic capacity (1.18x wild-type levels), and reduced glycolytic reserve (0.20x wild-type levels) as compared to wild-type cells (Fig. 1G). CTR1 KO respiration and glycolysis phenotypes correlated with modifications in energy charge (Extended Data Fig. 1-1F) (Atkinson and Walton, 1967) and lactate levels (Extended Data Fig. 1-1G), which were fully or partially restored by elesclomol (Extended Data Fig. 1-1). Together, these findings demonstrate that CTR1 KO cells have compromised copper-dependent Golgi and mitochondrial enzymes. The impairment of the function and organization of the respiratory chain in CTR1 KO cells induces a metabolic shift favoring glycolysis over oxidative phosphorylation.

### Unbiased discovery of copper deficiency mechanisms using proteomics and NanoString transcriptomics

Next, we sought to comprehensively identify pathways and signal transduction mechanisms altered by CTR1 KO copper deficiency. We performed quantitative mass spectrometry of the whole cell and phosphorylated proteomes by Tandem Mass Tagging (Fig. 2, source data in Extended Data File 1), focusing on the phosphoproteome as multiple kinases are known to have copper-binding domains which regulate their activity (Brady et al., 2014; Tsang et al., 2020). We quantified 8,986 proteins and 19,082 phosphopeptides (from 4066 proteins) across wild type cells and two CTR1 KO clones (Fig. 2A volcano plot, Extended Data File 1). The proteome and phosphoproteome did not segregate by genotype in principal component analysis until after thresholding data (p<0.01, fold change ≥1.5; Fig. 2B, PCA). This indicates discrete modifications of the global proteome and phosphoproteome in CTR1 KO cells. We identified 210 proteins and 224 phosphopeptides (from 162 proteins) that exhibited differential abundance in CTR1 KO cells (p<0.01, fold change ≥1.5, source data in Extended Data File 1), of which 26 proteins differed in both abundance and phosphorylation. 153 proteins and 138 phosphopeptides were more abundant while 57 proteins and 86 phosphopeptides decreased in abundance in CTR1 KO cells as compared to wildtype, respectively.

**Figure 2.**
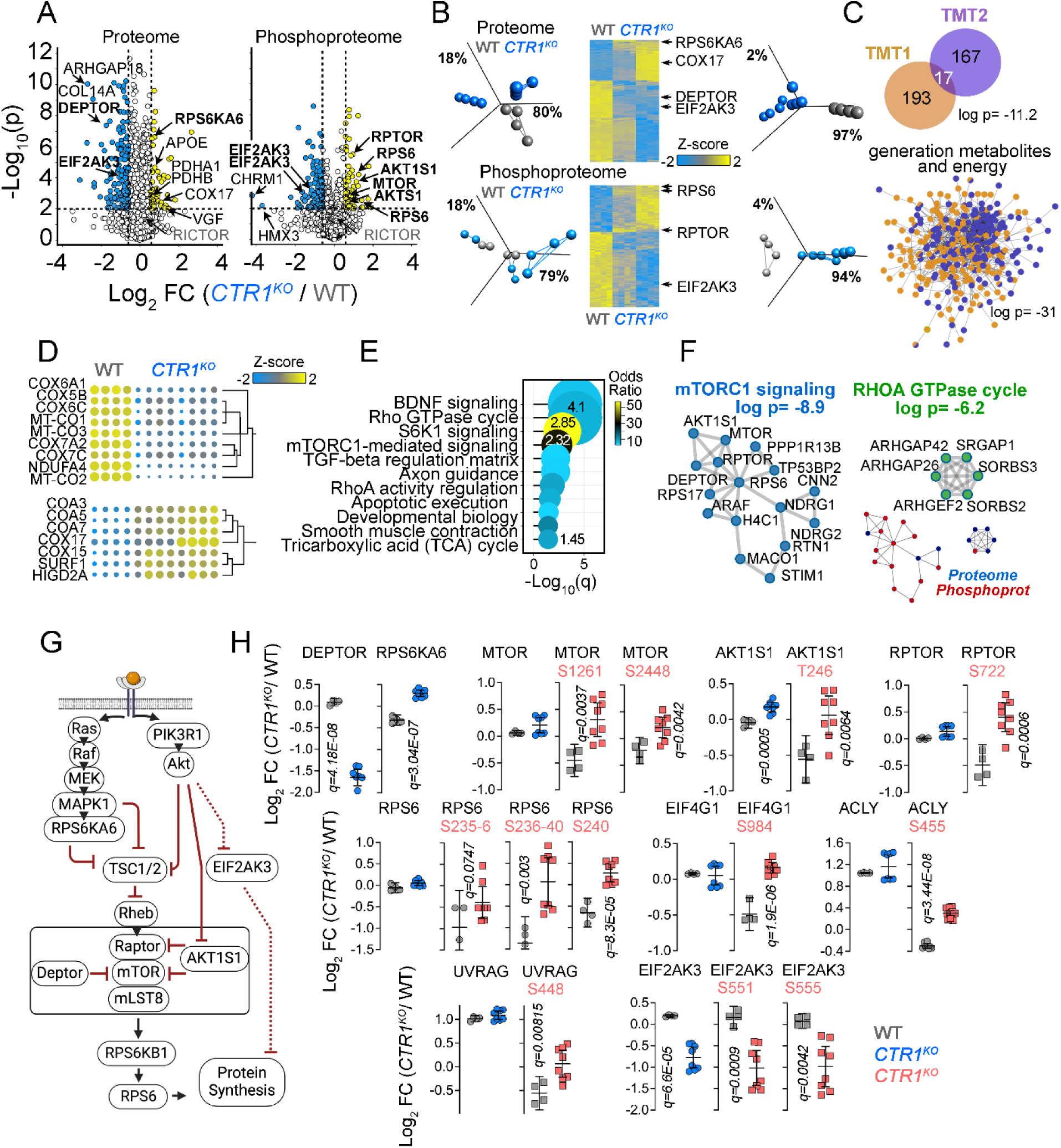
CTR1 mutant proteome and phosphoproteome have increased activation of mTOR-Raptor-S6K signaling and protein synthesis pathways. **A.** Volcano plots of the CTR1 KO cell proteome and phosphoproteome (TMT1), where yellow dots represent proteins or phosphoproteins whose expression is increased in KO cells and blue dots represent decreased expression in KO cells. n=4 for wild type cells and n=4 for KO cells in two independent clones (KO3 and KO11). **B.** Principle component analysis (PCA) of the whole proteome and phosphoproteome from wild type (gray) and two CTR1 KO clonal lines (blue symbols). Hierarchical clustering and PCA of all proteome or phosphoproteome hits where differential expression is significant with q<0.05 and a fold of change of 1.5 (t-test followed by Benjamini-Hochberg FDR correction). **C.** Replication TMT Proteome (TMT2) in independent CTR1 KO clone experiment (KO3 and KO20), Venn diagram overlap p value calculated with a hypergeometric test. Merged protein-protein interaction network of both TMT experiments is enriched in the GO term GO:0006091 generation of precursor metabolites and energy. **D.** TMT proteome levels of Complex IV subunits and assembly factors expressed as Z-score from TMT2. All 16 proteins changed more than 1.5-fold as compared to wild type cells with a q<0.05. **E.** Gene ontology analysis of differentially expressed proteins or phosphopeptides in CTR1 mutant proteome and phosphoproteome (TMT1). The Bioplanet database were queried with the ENRICHR engine. Fisher exact test followed by Benjamini-Hochberg correction. **F.** Metascape analysis of the proteome and phosphoproteome (TMT1). Ontology enrichment analysis was applied to a protein-protein interaction network of all components to select molecular complexes with MCODE based on significant ontologies. **G.** mTOR signaling pathway diagram modified from KEGG map04150. **H.** Mass spectrometry quantification of ontologically selected proteins and phosphopeptides (TMT1). Proteins are shown with blue circles and phosphopeptides are shown with red squares. Vertical numbers represent q values. See Extended Data File 1 for source data.

As further confirmation of Complex IV impairments and metabolic dysfunction in CTR1 KO cells (Fig. 1, source data in Extended Data Fig. 1-1), we used a replication TMT proteome (TMT2) to quantify Complex IV subunits and chaperones. Overall, we identified 184 differentially expressed proteins in CTR1 KO cells (Fig. 2C, Venn diagram, TMT2). These two proteome datasets overlapped 9.1 times above what is expected by chance (p=6.7E-12, hypergeometric probability). Both datasets are enriched 11-fold in a protein interaction network annotated to the GO term generation of precursor metabolites and energy (GO:0006091, p=1E-31, hypergeometric probability; Fig. 2C, Venn diagram and interactome, Extended Data File 1). This network of proteins included 9 proteins belonging to Complex IV, such as MT-CO1, MT-CO2, and MT-CO3, all of which were decreased in CTR1 KO cells ≥ 1.5-fold (p<0.01; Fig. 2D, Complex IV subunits, upper heat map, Extended Data File 1). This finding is consistent with the reduction of Complex IV subunits and supercomplexes by immunoblot and reduced respiration in CTR1 KO cells (Fig. 1C-F). This protein network also included seven Complex IV assembly factor and copper chaperones, such as COX17, whose levels were increased in CTR1 KO cells ≥ 1.5-fold (p<0.01; Fig. 2D Complex IV assembly factors, bottom heat map). Overall, this suggests copper-deficient cells upregulate assembly factors and chaperones for Complex IV in response to impaired assembly of this respiratory complex.

To identify ontological terms represented in the integrated CTR1 KO proteome and phosphoproteome (consisting of 345 total proteins with differential abundance and/or phosphorylation, source data in Extended Data File 1, TMT1), we queried the NCAST Bioplanet discovery resource with the ENRICHR tool and multiple databases with the Metascape tool (Huang et al., 2019). Both bioinformatic approaches identified the mTOR signaling pathway (ENRICHR q=4.76E-3 and Metascape q=9.8E-5), S6K1 signaling (ENRICHR q=1.4E-3), and Rho GTPase cycle in the top enriched terms (Fig. 2E, gene ontology, Extended Data File 1). Metascape analysis merged the CTR1 proteome and phosphoproteome into a network of 16 protein-protein interactions annotated to mTORC1-mediated signaling, which include these proteins: EIF4G1, mTOR, RPTOR (Raptor), AKT1S1, DEPTOR, RPS6, and RPS6KA6 (R-HSA-166208 and KEGG hsa04150, q=1.25E-9, z-score 22; Fig. 2F, mTORC1 and Fig. 2G, pathway diagram). As only 6 protein-protein interactions annotated to the Rho GTPase cycle (Fig. 2E-F, compare odds ratios), we focused on the mTOR-S6K pathway.

The most pronounced changes in the steady state mTOR proteome encompassed decreased levels of DEPTOR and increased levels of Ribosomal Protein S6 Kinase A6 (RPS6KA6 or RSK4) (Fig. 2H). DEPTOR is an mTOR inhibitor (Peterson et al., 2009), and RPS6KA6 is a member of the ribosomal S6K (RSK) family. RSK proteins (RSK1-RSK4) are activated downstream of MEK/ERK signaling, while the S6K family (S6K1 and S6K2) is phosphorylated by mTOR, but both families regulate translation through phosphorylation of ribosomal protein S6 (RPS6) and other related proteins (Meyuhas, 2015; Wright and Lannigan, 2023). These changes in abundance of DEPTOR and RPS6KA6 suggest heightened mTOR activity and RPS6 activation in copper deficiency. In support of this idea, CTR1 KO cells have increased phosphorylation of mTOR, RPS6, EIF4G1 (eukaryotic translation initiation factor 4 gamma 1), ACLY (ATP-citrate synthase), and UVRAG (UV radiation resistance associated) without changes in their steady state levels (Fig. 2H). The increased phosphorylation of these proteins was in phosphoresidues known to be responsive to mTOR activity, which include: mTOR S1261 and S2448, both correlated with increased mTOR activity and targets for the insulin/phosphatidylinositol 3–kinase (PI3K) pathway and the latter a target of RPS6KB1 (Ribosomal Protein S6 Kinase B1 or S6K1) (Chiang and Abraham, 2005; Holz and Blenis, 2005; Acosta-Jaquez et al., 2009; Copp et al., 2009; Rosner et al., 2010; Smolen et al., 2023); EIF4G1 S984 (Raught et al., 2000); RPS6 S235, S236, and S240, which are targets of S6K1 and RSK4 (Holz and Blenis, 2005; Roux et al., 2007; Magnuson et al., 2012; Meyuhas, 2015); and ACLY and UVRAG at S445 and S458, respectively (Kim et al., 2015; Covarrubias et al., 2016; Martinez Calejman et al., 2020). Increased phosphorylation of mTOR at S2448 and P70S6K at T389 was confirmed by immunoblot (Fig. 3C). We also observed increased protein expression and phosphorylation of AKT1S1 (PRAS40) at T246, a residue that is a target of AKT1 and whose phosphorylation correlates with active mTOR (Fig. 2H; Sancak et al., 2007). Importantly, we identified increased phosphorylation of RPTOR (Raptor, regulatory-associated protein of mTOR) at the RSK4 target residue S722 (Meyuhas, 2015) but observed no changes in RICTOR (RPTOR-independent companion of MTOR complex 2), suggesting that mTORC1 but not mTORC2 signaling is impacted (Ma and Blenis, 2009). Beyond the mTOR-S6K pathway, the proteome also revealed decreased protein levels of EIF2AK3 (PERK, eukaryotic translation initiation factor 2 alpha kinase 3), an ER stress response kinase and negative regulator of protein synthesis (Almeida et al., 2022), which was confirmed by immunoblot (Fig. 3C) and paralleled reduced content of two of its phosphopeptides (Fig. 2H, S551 and S555). While not known to be a direct target of mTOR, EIF2AK3 activity decreases by an AKT1-dependent mechanism (Peng et al., 2020). We conclude that genetic defects in CTR1 KO cells modify the proteome and signaling pathways converging on mTOR- and PERK-dependent signaling mechanisms upstream of protein synthesis pathways (Mounir et al., 2011; Hughes et al., 2020).

**Figure 3.**
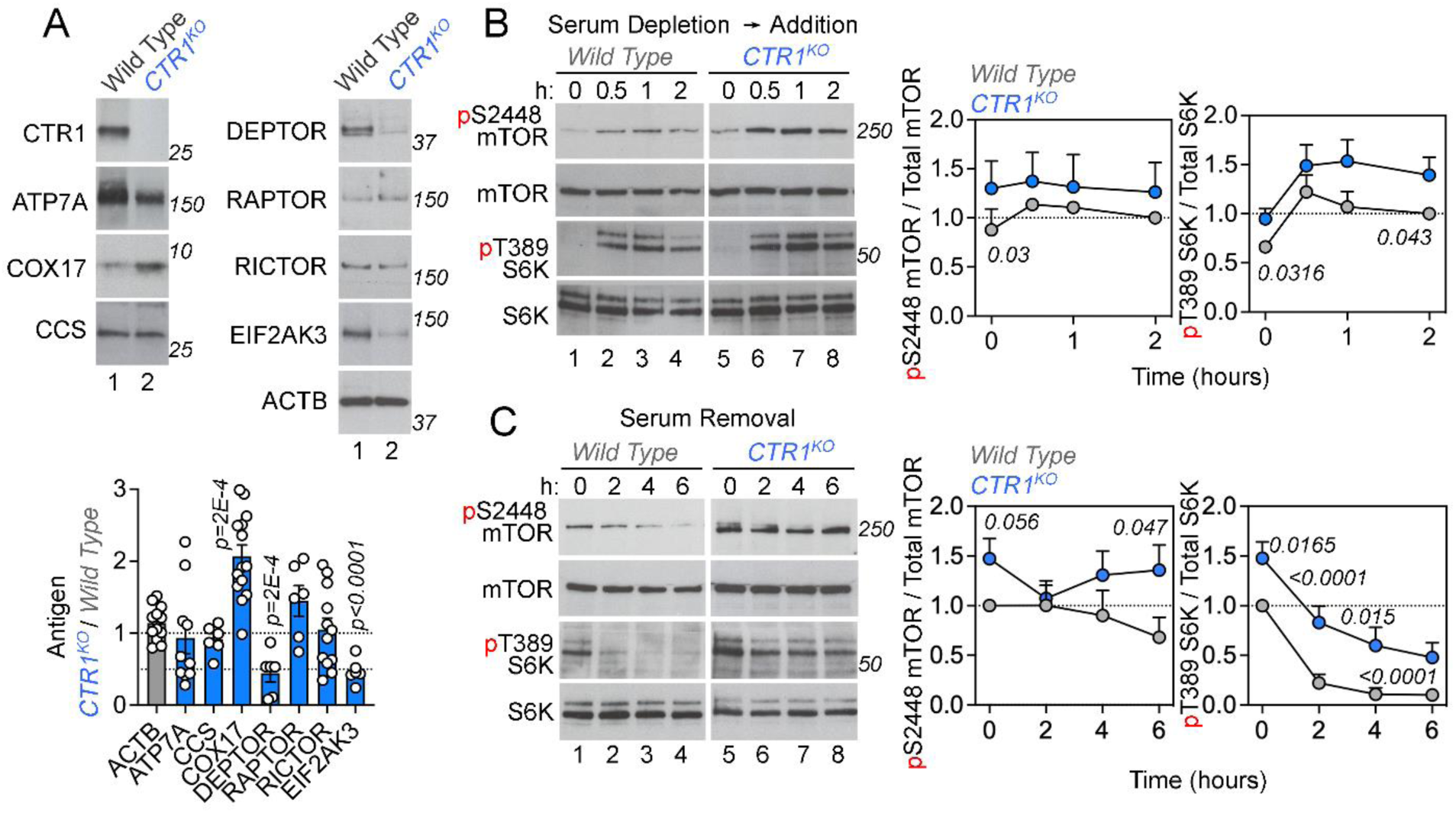
Increased activity of the mTOR-S6K pathway in CTR1 KO cells. **A.** Immunoblots of whole-cell extracts from wild-type and CTR1 mutant cells probed for ATP7A, COX17, CCS, DEPTOR, RAPTOR, RICTOR, and EIF2AK3 with actin as a loading control. Immunoblots were quantified by normalizing protein abundance to wild-type cells. Italicized numbers represent p values analyzed by two-sided permutation t-test. **B,C.** Immunoblots with antibodies detecting either phosphorylated or total mTOR or S6K as loading controls after overnight depletion of fetal bovine serum followed by serum addition for 0.5-2 hours (**B**, top) or at time 0 followed removal of fetal bovine serum for 2-6 hours (**C**, bottom). Graphs depict quantitation of blots on the left in 3-6 independent replicates as the ratio of the phosphorylated to total protein content, normalized to control at time 0 (**C**) or time at 2 hours (**B**) (Two-Way ANOVA followed by Benjamini, Krieger, and Yekutiel corrections).

We used NanoString nCounter transcriptomics as an orthogonal approach to proteomics to identify molecular mechanisms downstream of CTR1 KO-dependent copper deficiency. We examined steady-state mRNA levels using NanoString panels enriched in genes annotated to metabolic pathways and neuropathology processes, which collectively measure levels of over 1400 transcripts (Extended Data Fig. 2-1, Extended Data File 1), including 66 genes annotated to the mTOR pathway (KEGG). We identified 131 metabolic and 37 neuropathology annotated transcripts whose levels were altered in CTR1 KO cells (q<0.05 and fold of change ≥2, Extended Data Fig. 2-1A-C, Extended Data File 1). Metabolic transcripts were enriched in genes annotated to lysosome and mTOR signaling (KEGG, p<5.8E-6 and z-score >18, Extended Data Fig. 2-1D, Extended Data File 1). Similar ontology analysis but with the combined 168 transcripts whose levels were altered in CTR1 KO cells was enriched in genes annotated to central carbon metabolism in cancer and the PI3K-Akt signaling pathway (p=2.3E-11 z-score 8.8. and p=1.78E-6 and z-score 9.6, Extended Data File 1). With the exception of DEPTOR, a negative regulator of mTOR, 14 of the 15 transcripts annotated to mTOR and PI3K-Akt signaling were increased in CTR1 KO cells (Extended Data Fig. 2-1C). The copper dependency of these transcripts level changes was demonstrated using the copper chelator BCS. Differences in gene expression between wild type and CTR1 KO cells at baseline were magnified by BCS (Extended Data Fig. 2-1A,B, Extended Data File 1). A subset of these genes was sensitive to copper chelation in both wild-type and CTR1 KO cells, including PIK3R1, SLC7A5, and SLC3A2 (Extended Data Fig. 2-1B). Collectively, the metabolic transcriptome and proteome of CTR1 KO cells provide independent evidence of increased activation of PI3K-Akt and mTORC1-S6K signaling pathways.

### Increased steady-state activity of the mTOR-S6K pathway in CTR1 KO cells

To confirm the findings of our proteomic and transcriptomic datasets, we examined steady-state levels of several proteins identified in the CTR1 KO proteome by immunoblot and compared them to the housekeeping protein beta-actin (Fig. 3A). CTR1 KO cells exhibit increased levels of the copper chaperone COX17, as well as reduced levels of DEPTOR and EIF2AK3 (Fig. 3A). However, despite the severe copper depletion in CTR1 KO cells (Fig. 1B), we did not observe changes in the levels of ATP7A or CCS (Fig. 3A), two proteins frequently altered in copper depletion (Bertinato et al., 2003; Kim et al., 2010). In agreement with the proteomic profiling of these cells, there was no change in protein levels of RPTOR or RICTOR, both components of mTOR complexes (Fig. 3A).

We tested the hypothesis of heightened mTOR signaling in CTR1 KO cells by measuring the phosphorylation status of mTOR and p70/p85 S6K1. We used serum depletion and serum addition paradigms to inhibit or stimulate, respectively, mTOR-S6K and PI3K-Akt signaling pathway activity (Liu and Sabatini, 2020). We focused on mTOR S2448 and S6K1 T389 phosphorylation as sensors of mTOR signal transduction. mTOR S2448, which we identified in the CTR1 KO phosphoproteome (Fig. 2H), is present in the mTOR catalytic domain, is sensitive to nutrient availability and insulin, and is an S6K target (Navé et al., 1999; Reynolds et al., 2002; Cheng et al., 2004; Chiang and Abraham, 2005). The phosphoresidue T389 in S6K1 is phosphorylated by an insulin- and mTOR-dependent mechanism and indicates increased S6K activity (Burnett et al., 1998; Pullen et al., 1998; Liu and Sabatini, 2020). Relative to wild type cells, CTR1 KO cells had an increased mTOR and S6K1 phosphorylation at time 0 after an overnight serum depletion paradigm, revealing elevated mTOR activity even in the absence of an mTOR activating stimulus (Fig. 3B, compare lanes 1 and 5). Exposing these cells to serum progressively increased mTOR and S6K1 phosphorylation to a higher degree when comparing CTR1 KO to wild type cells (Fig. 3B, compare lanes 2-4 to 6-8). Additionally, CTR1 KO cells displayed increased phosphorylation of mTOR and S6K1 at baseline (0h in complete media) and over time after removal of serum (Fig. 3C), indicating that the mTOR signaling is resistant to serum deprivation in CTR1 KO cells. These results demonstrate that CTR1 KO cells have heightened activity of the mTOR-S6K signaling pathway.

### Activation of the mTOR-S6K signaling pathway is necessary for CTR1 KO cell survival

mTOR signaling is necessary for cell division, growth, and differentiation (Liu and Sabatini, 2020). We asked whether increased mTOR-S6K activation contributes to cell division and growth in CTR1 KO cells by measuring cell survival after pharmacological manipulation of mTOR activity. We reasoned that the increased mTOR signaling in CTR1 KO cells (Figs. 2, 3, Extended Data Fig. 2-1) would render mutant cells more sensitive to mTOR inhibition as compared to wild type cells. We quantitively assessed whether mTOR inhibition and cellular copper-modifying drugs interacted synergistically or antagonistically in cell survival assays (Ianevski et al., 2022). We used the zero-interaction potency (ZIP) model, which assumes there is no interaction between drugs, an outcome represented by a ZIP score of 0 (Yadav et al., 2015). ZIP scores above 10 indicate the interaction between two agents is likely to be synergistic, whereas a value less than -10 is likely to describe an antagonistic interaction (Yadav et al., 2015).

We first tested the individual effect of two activators of the mTOR-S6K signaling pathway, serum or insulin, on cell survival. CTR1 KO cells are more resistant to serum depletion and exhibit a reduced growth response when treated with insulin, which activates Akt and stimulates mTOR activity (Liu and Sabatini, 2020) (Fig. 4A). This is consistent with increased mTOR activation in these cells. We next asked whether serum and the mTOR inhibitors rapamycin and Torin-2 (Extended Data Fig. 4-1) would interact in a genotype-dependent manner to affect cell survival. Rapamycin is the canonical mTORC1 inhibitor, while Torin-2 inhibits both mTORC1 and mTORC2 with increased specificity and potency (Ballou and Lin, 2008; Zheng and Jiang, 2015). Based on our model, a synergistic response between serum and mTOR inhibitors (increased survival beyond the pro-survival effects of serum alone) would indicate that mTOR activity is deleterious for cell survival. Alternatively, an antagonistic response (negation of the pro-survival effect of serum by mTOR inhibition) would suggest cells are dependent on mTOR signaling for survival. Our results supported the latter, as the interaction between serum and either rapamycin or Torin-2 was antagonistic in both control and CTR1 KO cells (Fig. 4B-D, Extended Data Fig. 4-2A). The antagonism between serum and both rapamycin and Torin-2 was more pronounced in CTR1 KO as compared to wild type cells (ZIP score between -22.1 to -27.7 for wild type and -30.9 to -34.6 for CTR1 mutant cells, Fig. 4C), signifying that inhibiting mTOR is more detrimental to the pro-survival effects of serum in CTR1-null cells.

**Figure 4.**
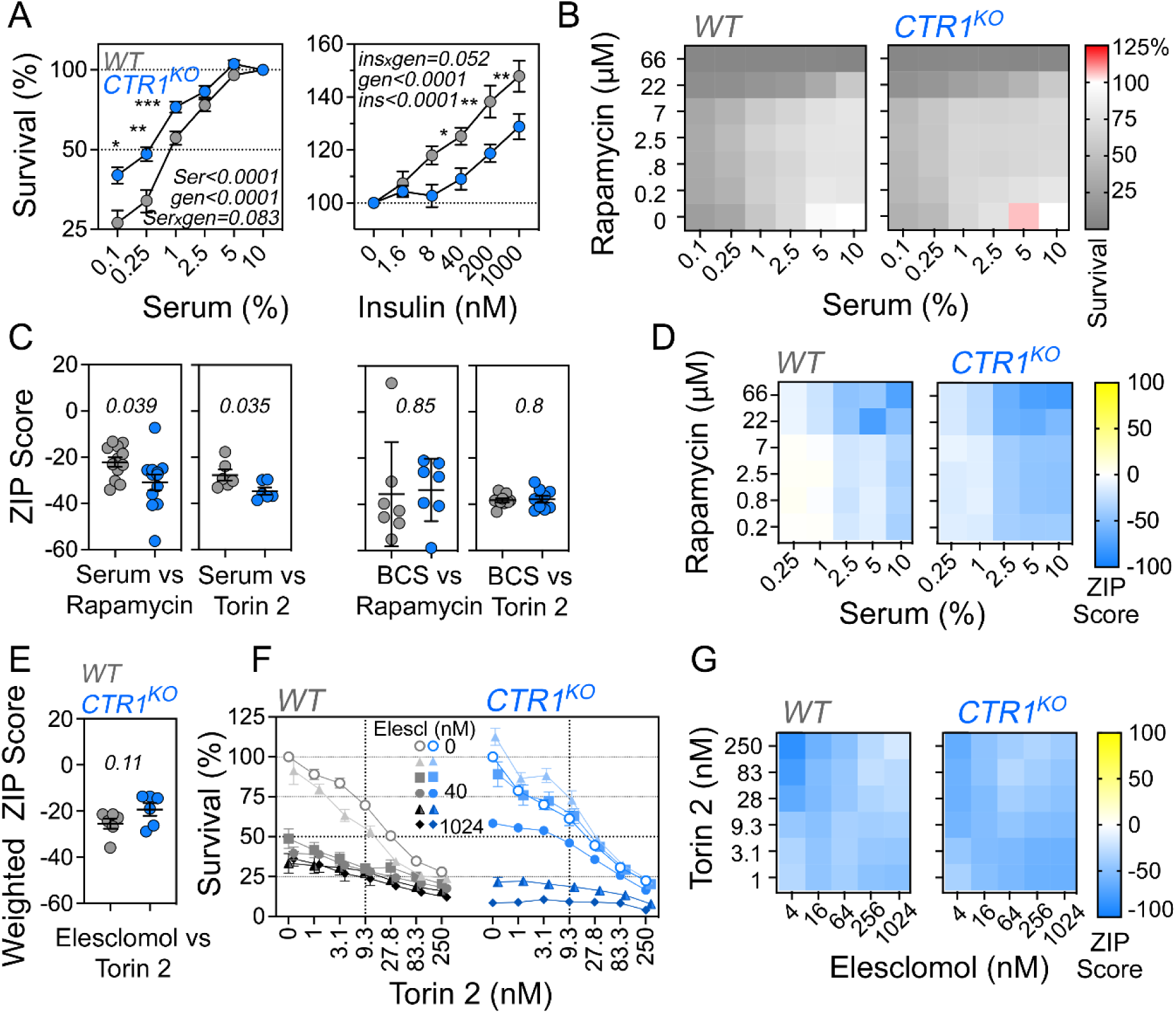
CTR1 knockout increases susceptibility to mTOR inhibition. **A.** Cell survival analysis of CTR1 mutants with increasing concentrations of serum or insulin (average ± SEM, n = 7 for serum and 5 for insulin, two-way ANOVA followed by Benjamini, Krieger, and Yekutieli corrections). **B-G.** Synergy analysis of cell survival of CTR1 mutants treated with increasing concentrations of combinations of the compounds serum, rapamycin, Torin-2, BCS, and elesclomol. **B.** Cell survival map for cells treated with serum and rapamycin, with the corresponding interaction synergy map calculated using the Zero Interaction Potency (ZIP) score for cell survival (Yadav et al., 2015) (**D**). **C-G.** Scores below -10 indicate an antagonistic interaction between the compounds. Maps were generated with at least six independent experiments per pair that generated percent cell survival maps presented in Extended Data Fig. 4-1 and average ZIP score for drug interactions in **C** or weighted ZIP score in **E** (see Methods). Average ± SEM, two-sided permutation t-test. **E-G.** Synergy analysis of CTR1 mutants with increasing concentrations of Torin-2 and elesclomol, with different colors and symbols indicating increasing concentrations of elesclomol (**F**) with average weighted ZIP score (**E**, two-sided permutation t-test) and elesclomol ZIP interaction synergy map (**G**).

Importantly, genotype-dependent differences in cell survival after mTOR inhibition were sensitive to pharmacological manipulation of copper levels. mTOR inhibitors in combination with the copper chelator BCS abrogated ZIP score differences between genotypes (-35.6 to -38.3 for wild-type and -33.8 to -37.9 for CTR1 KO, Fig. 4C, Extended Data Fig. 4-B,C). Conversely, low doses of elesclomol, which are sufficient to rescue copper content phenotypes and mitochondrial respiration in CTR1-null cells (Fig. 1B,E,F, Extended Data Fig. 1-1D,E), rendered CTR1 KO cells more resistant to increasing concentrations of Torin-2 (Fig. 4E-G, Extended Data Fig. 4-2D). These pharmacogenetic epistasis studies indicate that CTR1-null cells are more dependent on mTOR activity for their survival in a copper-dependent manner. These findings support a model where the activation of mTOR is an adaptive response in CTR1 mutant and copper-deficient cells.

### Interaction between mitochondrial respiration and increased protein synthesis in CTR1 KO cells

CTR1-null cells have increased activity of the mTOR-S6K pathway and decreased content of PERK (EIF2AK3), predicting increased protein synthesis in CTR1 KO cells as compared to wild type. We measured protein synthesis using puromycin pulse labeling of the proteome (Schmidt et al., 2009). Indeed, CTR1 KO cells display a 1.5-fold higher content of peptidyl-puromycin species as compared to wild type cells (Fig. 5A, Extended Data Fig. 5-1A-B). Puromycin incorporation was sensitive to the cytoplasmic protein synthesis inhibitor emetine in both genotypes (Fig. 5A). These results demonstrate increased protein synthesis in CTR1 mutant cells (Fig. 5A, Extended Data Fig. 5-1A-B). To further explore the mechanism by which mTOR-S6K activation and upregulation of protein synthesis may be an adaptive response by CTR1 KO cells, we tested whether cell survival and mitochondrial respiration were susceptible to protein synthesis inhibition in a genotype-dependent manner. We indirectly inhibited protein synthesis using serum depletion or directly with emetine (Fig 5A; Grollman, 1966; Mukhopadhyay et al., 2016). While CTR1 KO cell survival was resistant to serum depletion (Fig. 4A), we found a discrete yet significant decrease in cell survival in CTR1 KO cells after emetine addition (Extended Data Fig. 5-1C,D).

**Figure 5.**
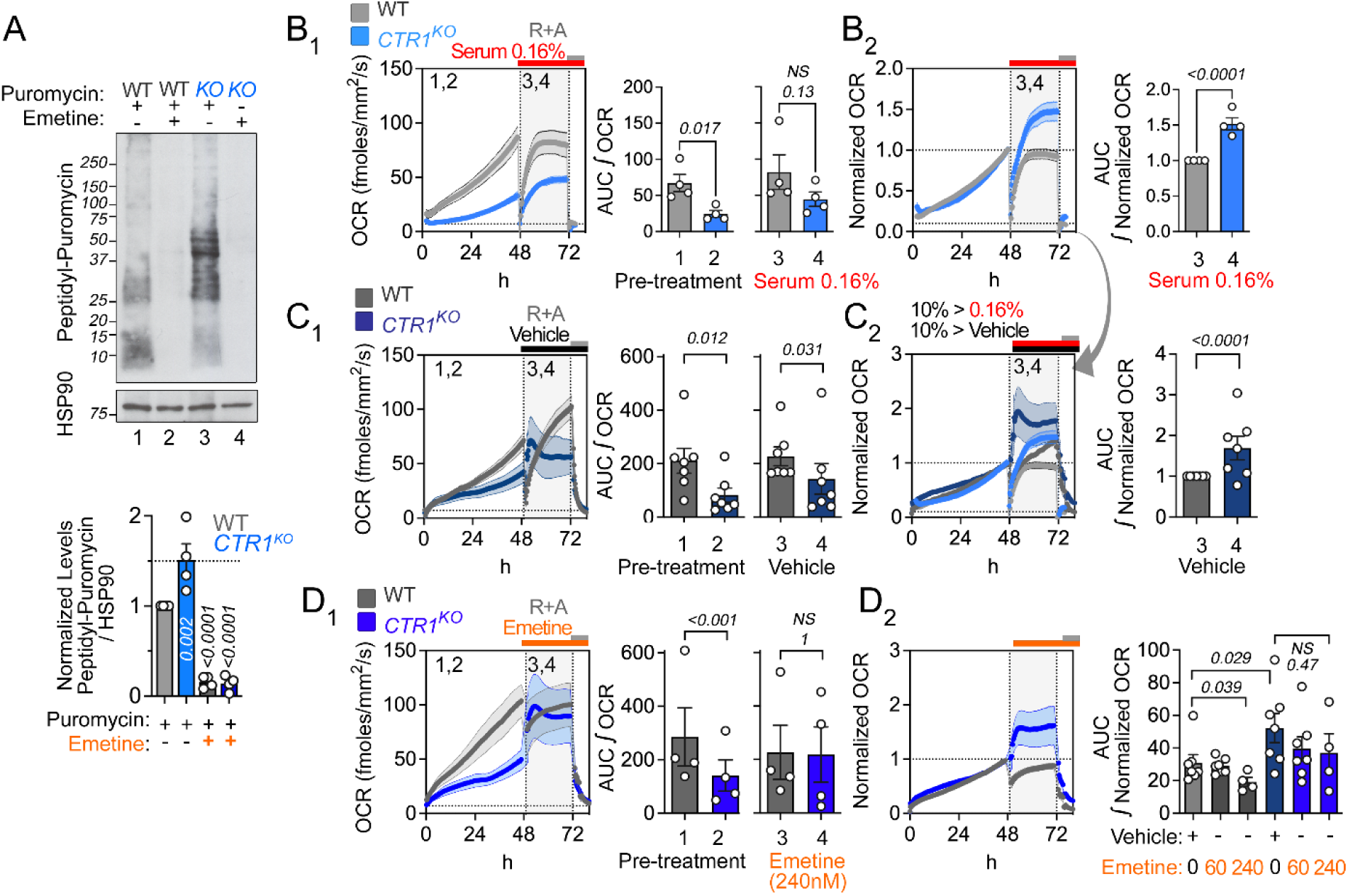
CTR1 mutant cells are resistant to protein synthesis inhibition. **A.** Immunoblot for puromycin in wild-type (lanes 1 and 2) and CTR1 mutant cells (lanes 3 and 4) treated with either vehicle (lanes 1 and 3) or 240 nM emetine (lanes 2 and 4) for 24 hours, followed by a 30 min pulse of puromycin. Quantification of the puromycin signal between 250 and 15 kDa normalized to HSP90. One-Way ANOVA, followed by Holm-Šídák’s multiple comparisons test. CTR1 clone KO20 was used for all experiments. **B-D.** Resipher respiration rates in wild type and CTR1 KO cells. Cells were grown in complete 10% serum media unless otherwise specified. Cells were incubated for 48h, followed by serum depletion (**B**, serum 0.16%), vehicle (**C**, DMSO), or emetine (**D**, 60 or 240 nM) for 24h. **B-D.** Assay was terminated at 72 h by the addition of rotenone plus antimycin (R+A). Columns 1 and 2 represent raw or normalized OCR, respectively, presented as OCR over time or the integrated area under the curve (AUC) for the indicated time periods. Each dot depicts a batch of concurrent experiments (n=4-7 per genotype for each experiment, average ± SEM, two-sided permutation t-test).

To measure whole-cell mitochondrial respiration (as defined by its abrogation with a mix of rotenone plus antimycin) over extended periods of time, we employed the Resipher system, which utilizes platinum organo-metallic oxygen sensors (Grist et al., 2010; Wit et al., 2023). In contrast with Seahorse, which requires serum-free media and measures oxygen consumption over a few hours, Resipher allows oximetry over prolonged periods while maintaining cells in their own milieu. We first measured respiration continuously over 48 h in standard media with 10% fetal bovine serum. Respiration increased over time in both genotypes, a reflection of cell number expansion (Fig. 5B1). CTR1 KO cell respiration was 35% of wild type levels (Fig. 5B1), a phenotype that cannot be explained by differences in cell numbers between genotypes (Extended Data Fig. 5-1E) and reproducing the respiratory phenotypes observed in Seahorse (Fig. 2E, compare columns 1 and 2, 53% of wild type basal respiration). However, a switch to low serum media decreased mitochondrial respiration in both genotypes (Fig. 5B1). To compare the relative effect of serum depletion on each genotype, we normalized the OCR values to the last time point before serum switch and revealed that CTR1-null cells respire 1.5 times more efficiently than wild type cells during serum depletion (Fig. 5B2). This effect cannot be explained by changes in cell number after switching to low serum (Extended Data Fig. 5-1E). These results show that mitochondrial respiration is resistant to serum depletion in CTR1-null cells.

Next, we compared respiration in CTR1 KO and wild type cells treated with fresh complete media containing either vehicle or emetine to directly inhibit cytoplasmic protein synthesis (Fig. 5C-D). First, we observed genotype-dependent effects of vehicle treatment. After the addition of new media, wild-type cells respire at the same rate as before the media switch (Fig. 5C), but CTR1 KO cells rapidly increase respiration, with a brief period where their raw OCR values are even greater than wild-type cells (Fig. 5C1). Relative to the last timepoint before media switch, CTR1 KO cells respire 1.7-fold more efficiently than wild-type cells (Fig. 5C2). Surprisingly, an increase in raw OCR was also observed in CTR1 KO cells after emetine treatment even above what was observed in vehicle-treated KO cells (Fig. 5D1, compare to 3,4 in C1). Normalized wild type cell respiration was inhibited by 40% after emetine treatment (Fig. 5D2). In contrast, the normalized respiration was resistant to emetine in CTR1-null cells (Fig. 5D2). These effects cannot be attributed to differences in cell number (Extended Data Fig. 5-1F). We conclude that CTR1-deficient cell respiration is resistant to direct inhibition of protein synthesis.

### Upregulation of protein synthesis machinery in a mouse model of copper deficiency

Next, we wanted to determine whether animal models of copper deficiency would corroborate our observations in CTR1 KO cells that copper depletion increases mTOR-S6K pathway activity and upregulates multiple pathways promoting protein synthesis. To address this question, we performed spatial transcriptomics of the cerebellum of *Atp7a^flx/Y^ :: Vil1^Cre/+^* mice, a conditional mouse model of Menkes disease. These mice, which lack the copper efflux transporter ATP7A in intestinal enterocytes, are unable to absorb dietary copper, depleting the brain of this metal and inducing subsequent pathology that phenocopies Menkes disease (Wang et al., 2012). We selected the cerebellum as it is known to be one of the earliest brain regions affected in Menkes disease, with Purkinje cells being particularly affected (Barnes et al., 2005; Gaier et al., 2013; Zlatic et al., 2015). To focus on mechanisms that could serve as an adaptive response to brain copper deficiency, we chose to study the transcriptome of the cerebellum at postnatal day 10, a presymptomatic timepoint before Purkinje cell death and the rapid onset of high mortality in this model and other Menkes mouse models (Wang et al., 2012; Guthrie et al., 2020; Yuan et al., 2022). As Menkes is a sex-linked disease, we focused our analysis on male mice.

We first confirmed that *Atp7a^flx/Y^ :: Vil1^Cre/+^* mice exhibit systemic copper depletion by ICP-MS as previously reported (Extended Data Fig. 6-2A; Wang et al., 2012). Next, we used the NanoString GeoMx Digital Spatial Profiler to generate anatomically resolved, quantitative, and PCR amplification-free transcriptomes (Decalf et al., 2019; Merritt et al., 2020; Zollinger et al., 2020) for two regions of interest (ROIs): the Purkinje cell and granular layers of the cerebellum (Fig. 6A). Within these ROIs, we segmented our analyses by GFAP immunoreactivity to enrich for areas of illumination (AOIs) containing Purkinje cells (GFAP-) or astrocytes in the granular layer (GFAP+) (Fig. 6B, Extended Data Fig. 6-1A). Using the GeoMx Mouse Whole Transcriptome Atlas (mWTA) and a mouse cerebellar cortex atlas (Kozareva et al., 2021), we quantified the transcriptome in these ROIs with a sequencing saturation above 80% (Extended Data Fig. 6-1B). After data normalization, we analyzed 19,963 genes across 58 AOIs from 4 animals of each genotype selected across different cerebellar folia. 17,453 genes were expressed in 10% of AOIs, and 10,290 genes were expressed in 50% of AOIs (Extended Data Fig. 6-1C,D). Segmentation by GFAP produced expected patterns of gene expression in Purkinje cell or granular layer AOIs, such that genes associated with Purkinje cells like *Calb1* and *Pcp2* were highly expressed in corresponding AOIs and minimally expressed in AOIs within the granular layer (66-fold enrichment of *Calb1* in Purkinje cell AOIs, Fig. 6C, Extended Data Fig. 6-1E, Extended Data Fig. 6-2B,C). Genes associated with granule cells and astrocytes were enriched in AOIs within the granular layer but had little expression in Purkinje cell AOIs (Extended Data Fig. 6-2B,C). Furthermore, the transcriptome of wild type Purkinje cells was enriched in genes annotated to MitoCarta3.0 knowledgebase ((Rath et al., 2021), Extended Data Fig. 6-2F), metabolic ontologies including oxidative phosphorylation (MSigDB p6.5E-43, Fisher exact test followed by Benjamini-Hochberg correction) and the mTOR pathway (MSigDB p2.4E-12) as compared to the granular layer (Extended Data Fig. 6-2E-GH).

**Figure 6.**
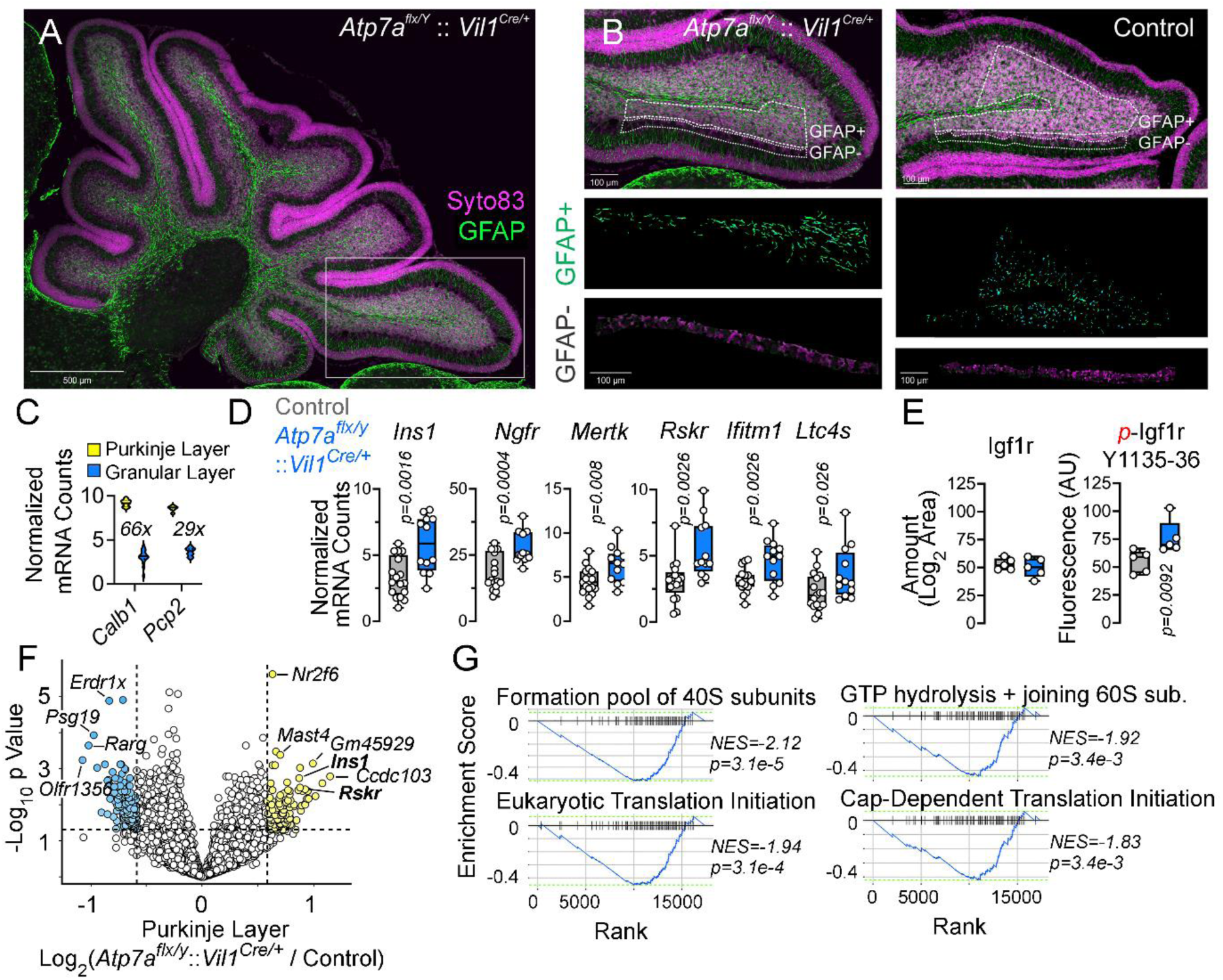
Transcriptome of cerebellar cortex layers in a presymptomatic Menkes mouse model. **A.** Sagittal sections of control and *Atp7a^flx/Y^ :: Vil1^Cre/+^* mutants at day 10 stained with Syto83 and GFAP to distinguish cerebellar layers. **B.** ROIs corresponding to the two AOIs analyzed: the GFAP-negative Purkinje cell layer AOI and the GFAP-positive granular layer AOI. **C.** Normalized mRNA counts for the Purkinje cell markers *Calb1* and *Pcp2* in the Purkinje and granular layer AOIs. **D.** Normalized mRNA counts for insulin and mTOR-S6 kinase related genes. Blue denotes mutant. N = 16 control AOIs and 13 mutant *Atp7a^flx/Y^ :: Vil1^Cre/+^*AOIs, 4 animals of each genotype. **E.** Abundance of Igf1r protein and p-Igf1r (Tyr1135/Tyr1136) as measured by LFQ-MS or Luminex, respectively. Box plots are all two-sided permutation t-tests. **F.** Volcano plot of mRNAs differentially expressed in mutant *Atp7a^flx/Y^ :: Vil1^Cre/+^* Purkinje neurons vs control. Yellow symbols mark genes with increased expression in mutant *Atp7a^flx/Y^ :: Vil1^Cre/+^*Purkinje cells. **G.** Gene Set Enrichment Analysis and normalized enrichment score (NES) of genes differentially expressed by comparing controls to mutant *Atp7a^flx/Y^ :: Vil1^Cre/+^* Purkinje cells. Gene sets enriched in mutants correspond to negative NES. p values are corrected. See Extended Data File 2 for raw data

In Purkinje cells from mutant mice, we revealed increased expression of several genes annotated to the Human MSigDB mTOR pathway (Liberzon et al., 2011). These include the most upstream mTOR activator, insulin (*Ins*, 1.75 fold); S6-related kinase (*Rskr*, 1.75 fold); Nerve Growth Factor Receptor (*Ngfr*, 1.48 fold); *Mertk*, a tyrosine kinase (1.42 fold); interferon-induced transmembrane protein 1 (*Ifitm1*, 1.5 fold); and leukotriene C4 synthase (*Ltc4s*, 1.66 fold) (Fig. 6D). At the protein level, we observed increased phosphorylation of the insulin-like growth factor 1 receptor Igf1r at Tyr1135/Tyr1136 with no change in expression of the receptor, as measured by Luminex or label-free mass spectrometry of the cerebellum, respectively (Fig. 6E). Notably, IGF1-R stimulation has been reported to activate the p70 S6K1 pathway (Cai et al., 2017). Together, this shows that brain copper deficiency increases the expression of components of the mTOR-S6K pathway activity in Purkinje cells. There were no genotype-dependent changes in expression of components of the electron transport chain in either the Purkinje or granular layer ROIs (Extended Data Fig. 6-3). We analyzed the spatial transcriptomic data by Gene Set Enrichment Analysis (GSEA) (Subramanian et al., 2005; Korotkevich et al., 2019) to unbiasedly identify pathways disrupted by brain copper depletion in cerebellar cortex. We observed a global upregulation of the protein synthesis machinery, including 82 ribosomal subunits and protein synthesis elongation factors, in Purkinje cells but not in the granular layer (Fig. 6G, FGSEA estimated p value Benjamini-Hochberg-correction). Expression levels of cytoplasmic ribosome transcripts were similar in wild type Purkinje cells and granular layer cells (Extended Data Fig. 6-2D). Our data support a model of mTOR-dependent upregulation of protein synthesis machinery in Purkinje neurons in presymptomatic copper-deficient mice.

### Genetic modulation of mTOR pathway-dependent protein synthesis activity modifies copper deficiency phenotypes in *Drosophila*

We genetically tested whether increased activity of mTOR pathway-dependent protein synthesis was an adaptive or maladaptive mechanism in copper deficiency. We studied the effect of mTOR-S6K pathway gain- and loss-of-function on copper-deficiency phenotypes in *Drosophila.* In animals overexpressing *ATP7*, which induces copper depletion by metal efflux (*ATP7*-OE, Figs. 7 and Extended Data Fig. 7-1; Norgate et al., 2006; Binks et al., 2010; Hwang et al., 2014; Gokhale et al., 2016; Hartwig et al., 2020), we also over-expressed and/or knocked down members of the mTOR pathway in either the epidermal epithelium (pnr-GAL4) or class IV sensory neurons (ppk-GAL4) (Fig. 7, Extended Data Fig. 7-1, Table 3; Binks et al., 2010; Gokhale et al., 2016; Hartwig et al., 2020). *ATP7* overexpression in the dorsal midline caused depigmentation and bristle alterations in both males and female (Extended Data Fig. 7-1; Binks et al., 2010; Gokhale et al., 2016; Hartwig et al., 2020). Loss of function of either S6k, raptor, or Akt by RNAi intensified epidermal *ATP7*-OE copper deficiency phenotypes (Extended Data Fig. 7-1), including the induction of additional dorsal thoracic caving (*RAPTOR*-IR and *AKT*-IR), necrotic tissue (*RAPTOR*-IR and *AKT*-IR), thoracic “dimples’ (*S6K*-IR in males, see arrowheads), and ultimately with increased lethality in response to *Akt* RNAi, with the few surviving animals showing caving and necrosis of the thoracic dorsal midline and/or scutellum (Extended Data Fig. 7-1). We did not detect overt *ATP7*-OE phenotype modifications by mTOR pathway transgene overexpression (*Akt* or *S6k*) or the two RNAi lines for *rictor* (Extended Data Fig. 7-1). These results suggest an adaptive role of mTOR-Raptor-S6K-dependent protein synthesis machinery in copper deficiency.

**Fig. 7.**
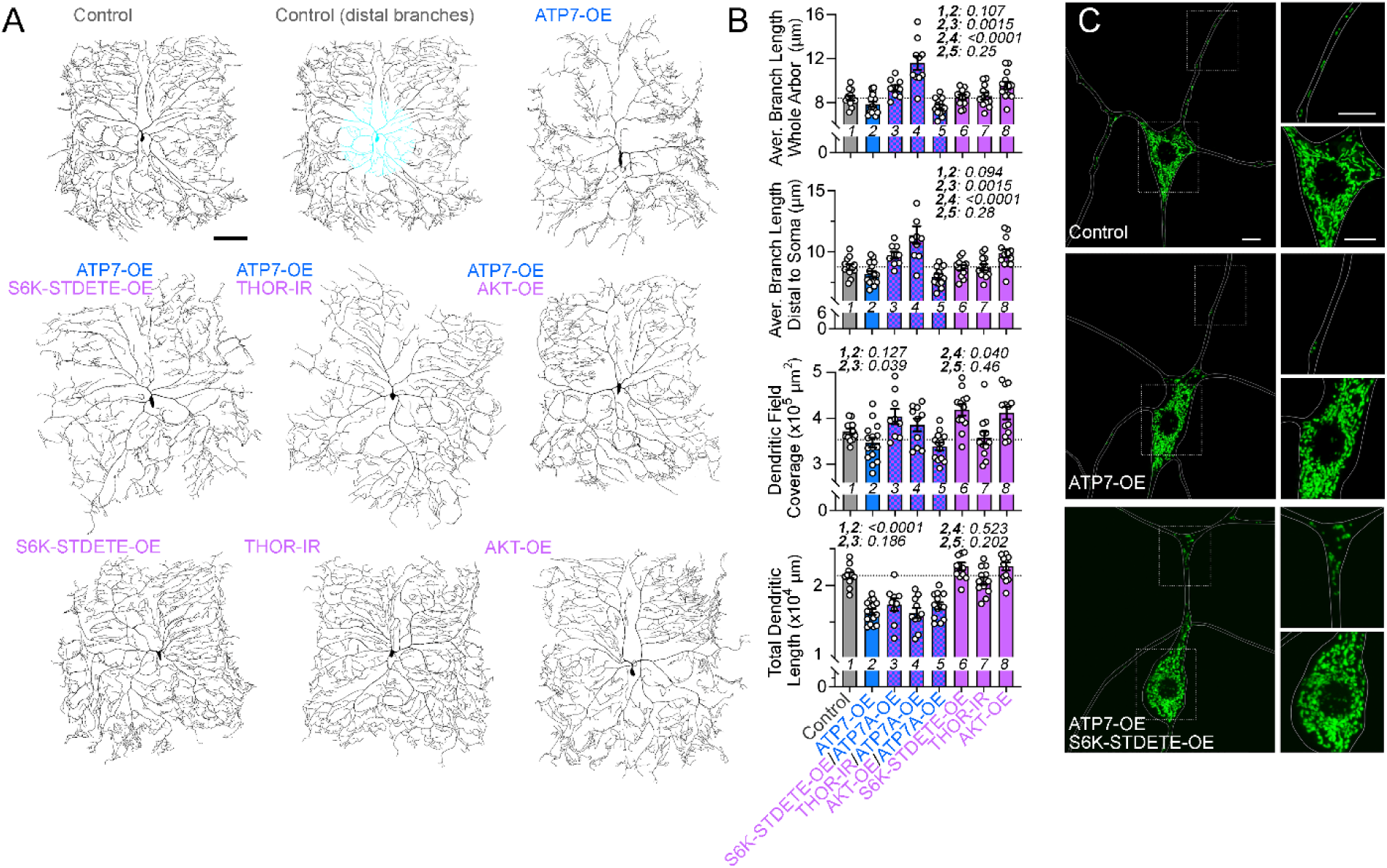
mTOR-dependent protein synthesis pathways ameliorate copper-depletion phenotypes in sensory neurons. **A.** Representative reconstructed dendritic arbors from live confocal images of C-IV da neurons of the specified genotypes labeled by GFP (see Table 3 and Methods.) Scale bar: 100 µm. Right panel depicts 400 μM circle used to distinguish proximal and distal dendrites (see Methods). **B.** Quantitative analysis of dendritic parameters in the specified genotypes: average branch length for the entire dendritic arbor or for the region distal to the soma (see Methods), dendritic field coverage, and total dendritic length. Each dot represents an independent animal. Average ± SEM. Italicized numbers represent q values (One-Way ANOVA, followed by Benjamini, Krieger, and Yekutieli multiple comparisons correction). **C.** Representative live confocal images of C-IV da neurons of the specified genotypes expressing a mitochondria-targeted GFP that were manually traced using a plasma membrane marker (CD4-tdTomato, not shown). Scale bar: 5 µm.

To test whether gain-of-function of mTOR-dependent protein synthesis machinery would rescue *ATP7*-OE phenotypes, we measured the complexity of class IV sensory neuron dendritic arbors and mitochondrial distribution in the third instar *Drosophila* larva. Dendrites in this cell type are a sensitive and quantitative reporter to measure cell-autonomous mechanisms in neuronal copper homeostasis (Fig. 7; Gokhale et al., 2016; Hartwig et al., 2020). Copper deficiency due to overexpression of *ATP7* in these neurons decreased the complexity of the dendritic arbor (as quantified by average branch length and total dendritic length; Fig. 7A-B) and depleted mitochondria from dendrites (Fig. 7C), as previously reported (Gokhale et al., 2016; Hartwig et al., 2020). The sole overexpression of *Akt* was not sufficient to modify *ATP7*-OE phenotypes (Fig. 7A-B). However, dendritic branch phenotypes were partially rescued by overexpression of either a constitutively active phosphomimetic mutant of *S6k* (*S6k*-STDETE, Fig. 7A-B; Barcelo and Stewart, 2002) or RNAi against *Thor*, the *Drosophila* orthologue of 4-EBP1, an inhibitor of protein synthesis downstream of mTOR necessary for mitochondrial biogenesis (Fig. 7A-B; Miron et al., 2001; Qin et al., 2016). 4E-BP1-dependent inhibition of protein synthesis is inhibited by mTOR-dependent phosphorylation; thus, 4-EBP1/Thor removal increases protein synthesis (Morita et al., 2013; Morita et al., 2015). *S6k*-STDETE-OE or *THOR*-IR increased the length of dendritic branches, a phenotype specific to the distal dendritic branches without changes in total dendritic length (Fig. 7A-B). This increase in the average branch length resulted in an expansion of the dendritic field coverage by class IV neurons (Fig. 7A-B). Simultaneously, mitochondria distribution was rescued in *ATP7*-OE neurons expressing *S6k*-STDETE-OE, with the redistribution of mitochondria to dendrites (Fig. 7C, not quantified). These results demonstrate that activation of mTOR-dependent protein synthesis mechanisms partially revert copper-depletion neuronal phenotypes. We conclude that mTOR-dependent protein synthesis is an adaptive mechanism in neuronal copper deficiency.

## Discussion

Here, we report that concomitant to changes in bioenergetics, copper-depleted cells undergo mTOR and PERK signaling modifications which converge on an adaptive response of increased protein synthesis in cellular and animal models of copper depletion.

We first established a cellular model of copper depletion by CRISPR editing to knockout CTR1 in SH-SY5Y cells (Fig. 1, Extended Data Fig. 1-1). This cellular system recapitulates cardinal phenotypes of copper depletion such as compromises in copper-dependent enzymes including Complex IV of the respiratory chain, which drives cells into glycolysis with increased cellular lactate content (Fig. 1, Extended Data Fig. 1-1). This is consistent with previous reports that of the respiratory chain complexes, complex IV specifically is impaired in copper deficiency (Zeng et al., 2007; Ghosh et al., 2014; Soma et al., 2018). Importantly, respiration and metabolic phenotypes observed in CTR1 KO cells can be reverted by elesclomol (Fig. 1, Extended Data Fig. 1-1), a drug that rescues neurodegeneration and organismal survival in animal models of copper depletion (Guthrie et al., 2020; Yuan et al., 2022).

We used the CTR1 KO cell model to unbiasedly identify cellular processes sensitive to copper depletion. The proteome, phosphoproteome, and metabolic transcriptome of CTR1-null cells identified the mTOR signaling pathway as one of the most enriched terms (Fig. 2, Extended Data Fig. 2-1). Patterns of differential gene, protein, and phosphoresidues levels are indicative of increased mTORC1-S6K activation and downstream activation of protein synthesis through increased levels and phosphorylation of RPS6 (Fig. 2, Fig. 3). At least one of the mechanisms of this mTOR activation is through decreased expression and protein levels of the mTOR inhibitor DEPTOR (Fig. 2, Extended Data Fig. 2-1, Fig. 3). We suspect additional mechanisms, such as increased receptor tyrosine activity, also contribute to increased mTOR activation, as evidenced by increased insulin-dependent activity observed in Purkinje neurons of copper-deficient *Atp7a^flx/Y^ :: Vil1^Cre/+^* mice (Fig. 6) and by the effect of *Akt*, *raptor*, and *S6k* RNAi enhancing *Drosophila* copper depletion epidermal phenotypes (Extended Data Fig. 7-1). The effects mTOR activation in CTR1 KO cells span the RPTOR/Raptor (mTORC1) branch of the pathway, including increased expression of 14 transcripts annotated to mTOR and/or PI3K-Akt signaling (Extended Data Fig. 2-1). Moreover, numerous proteins annotated to mTORC1-mediated signaling pathway exhibited increases in phosphorylation at residues responsive to mTOR and/or S6K activity, including mTOR, RPS6, EIF4G1, ACLY, UVRAG, and AKT1S1 (Fig. 2). In support of the idea that CTR1-null cells increase mTORC1 signaling and downstream S6K activation, these cells exhibit increased phosphorylation at mTOR S2448 and S6K1 T389 at baseline as well as under conditions of serum depletion or addition as compared to wild-type cells (Fig. 3). A notable finding is the discovery that concomitant to an upregulation of mTOR signaling and activation of S6K, we observed decreased protein expression and phosphorylation of EIF2AK3 (PERK) (Fig. 2, Fig. 3). As is expected by these changes in PERK and mTORC1 signaling, CTR1-null cells upregulate protein synthesis over wild-type cells (Fig. 5, Extended Data Fig. 5-1). Similarly, *Atp7a^flx/Y^ :: Vil1^Cre/+^* mice upregulate the protein synthesis machinery and expression of several genes in the mTOR pathway in Purkinje neurons at an early timepoint before the onset of cell death and mortality (Fig. 6; Wang et al., 2012; Guthrie et al., 2020; Yuan et al., 2022).

Increased mTOR signaling and protein synthesis in response to copper depletion appears to be a cell-type specific response that is observed in human SH-SY5Y neuroblastoma cells (Fig. 2-5) and Purkinje cells but not granular layer cells in *Atp7a^flx/Y^ :: Vil1^Cre/+^* mice (Fig. 6), suggesting that Purkinje cells are especially sensitive to copper depletion. CTR1 KO mouse embryonic fibroblasts were previously reported to show no changes in phosphorylation of either mTOR- or PI3K-Akt-specific substrates (Tsang et al., 2020). In Menkes disease, Purkinje cells experience rapid and specific pathology relative to other cell types and regions of the brain, including enlarged and distended mitochondria, increased dendritic arborization, and cell death, which have been reported in both humans (Ghatak et al., 1972; Vagn-Hansen et al., 1973; Purpura et al., 1976; Hirano et al., 1977; Troost et al., 1982; Kodama et al., 2012) and various mouse models (Yamano and Suzuki, 1985; Niciu et al., 2007; Lenartowicz et al., 2015; Guthrie et al., 2020). Importantly, some of these phenotypes are recapitulated by cell-autonomous hyperactivation of mTOR; however, in contrast with Menkes disease, mTOR activation in copper-sufficient Purkinje cells *increases* the size and respiratory activity of their mitochondria (Sakai et al., 2019). Dramatic differences in the mitochondrial proteome between Purkinje cells and other cell types in the cerebellum have been reported (Fecher et al., 2019). Thus, Purkinje neurons and their mitochondria may be uniquely susceptible to copper depletion due to cell-type specific mitochondrial properties and/or abundance, as well as their dependency on mTOR signaling (Extended Data Fig. 6-2). This is consistent with the enrichment of nuclear-encoded mitochondrial transcripts and mitochondrial activity in GABAergic neurons, particularly those expressing parvalbumin (Wynne et al., 2021; Bredvik and Ryan, 2024).

Previous reports have connected mitochondrial dysfunction with mTOR or PERK signaling. Mouse models of Leigh syndrome (caused by a deficiency in the Complex I subunit Ndufs4) and mitochondrial myopathy exhibit increased mTORC1 activity (Johnson et al., 2013; Khan et al., 2017), and downregulation of the mitochondrial respiratory complexes I, III, or IV stimulates TOR activity in the *Drosophila* wing disc (Perez-Gomez et al., 2020). Multiple diseases with impairments in mitochondrial respiration have been reported to benefit from mTOR inhibition, which is proposed to help alleviate metabolic stress. For example, rapamycin promotes survival and ameliorates pathology in a mouse model of Leigh syndrome (Johnson et al., 2013), and death due to energy stress in cells that are Coenzyme-Q-deficient is rescued by several mTORC1/2 inhibitors as well as protein synthesis inhibition by cycloheximide (Wang and Hekimi, 2021). These studies stand in contrast with our results. Our pharmacogenetic epistasis studies in CTR1 KO cells demonstrate that mTOR activation and increased protein synthesis is adaptive (Fig. 4, Extended Data Fig. 4-2) and that these cells are more sensitive to protein synthesis inhibition (Fig. 5, Extended Data Fig. 5-1). Additionally, stimulating protein synthesis in copper depleted *Drosophila* by either *S6k* overexpression or *Thor* RNAi partially rescues dendritic branching and mitochondrial phenotypes in class IV sensory neurons (Fig. 7), while downregulation of *S6k*, *Akt*, or *raptor* increases the severity of copper deficiency phenotypes in the epidermis (Extended Data Fig. 7-1). This suggests that mTOR-Raptor-S6K-dependent protein synthesis is adaptive in copper-deficient neurons and is necessary and sufficient to partially revert the effects of copper depletion in *Drosophila* (Fig. 7 and Extended Data Fig. 7-1). Based on these data and the fact that two distinct pathways favoring increased protein synthesis are activated in response to copper deficiency, we conclude that increased protein synthesis downstream of increased mTORC1 activity and/or decreased PERK represents a pro-survival response to copper depletion.

It is perhaps counterintuitive for cells to upregulate a nutrient sensing pathway like mTOR when deficient in an important micronutrient and enzyme cofactor like copper, which might be expected to decrease mTOR activity (Dennis et al., 2001). It is particularly surprising given that metabolic and mitochondrial diseases benefit from mTOR inhibition (see above). We speculate this could be a way to rectify an imbalance in proteostasis and metabolism due to copper depletion. Impaired autophagic flux has been reported in hyperglycolytic neurons (Jimenez-Blasco et al., 2024), and as copper is required for ULK1 activity and autophagy (Tsang et al., 2020), copper deficiency may inhibit autophagy, limiting the availability of a recycled pool of amino acids for *de novo* protein synthesis or as fuel for mitochondrial respiration. Thus, mTOR activation could lead to an increase in the efficiency of protein synthesis and cell cycle progression by increasing amino acid uptake from the media, as suggested by our findings of increased mRNA for the amino acid transporters SLC7A5 and SLC3A2 in CTR1 KO cells and wild-type cells treated with the copper chelator BCS (Extended Data Fig. 2-1), and/or by increased translation efficiency of particular RNA splice variants (Ma et al., 2008; Ma and Blenis, 2009). Increased levels of amino acid transporters at the cell surface may relate to the perplexing result that CTR1-null cells increase normalized respiration more than wild-type cells after the addition of fresh media (Fig. 5). While CTR1 KO cells exhibit increased glycolysis and lactate levels and decreased basal and ATP-dependent respiration under baseline condition (Figs. 1, 1-1), mTOR activation and the upregulation of COX17 and other chaperones for Complex IV may prime these cells to utilize nutrients such that fresh media enables increased respiration even under low serum conditions (Fig. 5C2, compare the blue groups) or when treated with emetine to inhibit protein synthesis (Fig. 5D2).

To our knowledge, our cellular and animal models of copper depletion provide the first evidence that 1) genetic defects that impair cellular copper homeostasis simultaneously modify two signaling pathways regulating protein synthesis and 2) in which the upregulation of protein synthesis is adaptive for cell-autonomous disease phenotypes. We propose that neuronal cell pathology occurs when resilience mechanisms engaged in response to copper deficiency are outpaced by the increasing bioenergetic demands of the cell during neurodevelopment.

## Materials and Methods

### Cell lines, gene editing, and culture conditions

Human neuroblastoma SH-SY5Y cells (ATCC, CRL-2266; RRID:CVCL_0019) were grown in DMEM media (Corning, 10-013) containing 10% FBS (VWR, 97068-085) at 37°C in 10% CO_2_, unless otherwise indicated. SH-SY5Y cells deficient in SLC31A1 were generated by genome editing using gRNA and Cas9 preassembled complexes by Synthego with a knock-out efficiency of 97%. The gRNAs used were UUGGUGAUCAAUACAGCUGG, which targeted transcript ENST00000374212.5 exon 3. Wild-type and mutant cells were cloned by limited dilution and mutagenesis was confirmed by Sanger sequencing with the primer: 5’GGTGGGGGCCTAGTAGAATA. All controls represent either a single wild-type clone or a combination of two wild-type clones. All experiments used two separate mutant clones of cells (KO3 and KO20, see Extended Data Fig. 1-1A,B) were used to exclude clonal or off-target effects unless otherwise indicated.

### Mouse husbandry

Animal husbandry and euthanasia was carried out as approved by the Emory University Institutional Animal Care and Use Committees. Genotyping was performed by Transnetyx using real-time PCR with the Vil1-Cre-1 Tg, Atp7a-2 WT, and Atp7a-2 FL probes.

### Antibodies

Table 1 lists the antibodies used at the indicated concentrations for western blots and immunofluorescence.

**Table 1.** Antibodies.

| **Antibody** | **Dilution** | **Catalog Number** | **RRID** |
| --- | --- | --- | --- |
| Actin B | 1:5000 | Sigma-Aldrich A5441 | AB_476744 |
| ATP7A | 1:500 | NeuroMab 75-142 | AB_10672736 |
| CCS | 1:500 | ProteinTech 22802-1-AP | AB_2879172 |
| COX17 | 1:500 | ProteinTech 11464-1-AP | AB_2085109 |
| COX4 | 1:1000 | Cell Signaling 4850 | AB_2085424 |
| CTR1 (SLC31A1) | 1:2000 | ProteinTech 67221-1-IG | AB_2919440 |
| DBH | 1:500 | Millipore AB1536 | AB_2089474 |
| DEPTOR | 1:1000 | Cell Signaling 11816 | AB_2750575 |
| GFAP-AF594 | 1:400 | Cell Signaling 8152 | AB_10998775 |
| HSP90 | 1:1000 | BD Biosciences 610418 | AB_397798 |
| mTOR | 1:1000 | Cell Signaling 2983 | AB_2105622 |
| mTOR pSer2448 | 1:1000 | Cell Signaling 5536 | AB_10691552 |
| NDUFB11 | 1:500 | Abcam ab183716 | AB_2927481 |
| OXPHOS mix | 1:250 | Abcam ab110412 | AB_2847807 |
| PERK (EIF2AK3) | 1:1000 | Cell Signaling 5683 | AB_10841299 |
| Puromycin | 1:500 | Sigma MABE342 | AB_2737590 |
| Raptor | 1:1000 | Cell Signaling 2280 | AB_561245 |
| Rictor | 1:1000 | Cell Signaling 2114 | AB_2179963 |
| S6K P70 | 1:1000 | Cell Signaling 9202 | AB_331676 |
| S6K pThr389 | 1:1000 | Cell Signaling 9234 | AB_2269803 |
| SDHA | 1:1000 | 11998 | AB_2750900 |
| UQCRC2 | 1:500 | Abcam ab14745 | AB_2213640 |
| Mouse IRDye 680RD | 1:1000 | LI-COR 926-68070 | AB_10956588 |
| mouse HRP | 1:5000 | A10668 | AB_2534058 |
| rabbit HRP | 1:5000 | G21234 | AB_2536530 |

### Drugs

Table 2 lists the drugs used at the indicated concentrations or concentration ranges as described in their corresponding figure legends.

**Table 2.** Drugs.

| **Drug** | **Source/Cat. No** | **Storage/Stock** | **Concentration range** |
| --- | --- | --- | --- |
| Elesclomol | VWR, 101758-608 | 1 mM, DMSO, -20C | 1 nM - 1024 nM |
| Copper chloride | Sigma, 203149 | 120 mM, water, -20C | 25 μM - 600 μM |
| BCS | Sigma, B1125 | 400 mM, DMSO, -20C | 0.1 mM - 1.6 mM |
| Oligomycin | Sigma, 75351 | 10 mM, DMSO, -20C | 1.0 μM |
| FCCP | Sigma, C2920 | 10 mM, DMSO, -20C | 0.25 μM |
| Rotenone | Sigma, R8875 | 10 mM, DMSO, -20C | 0.5 μM |
| Antimycin A | Sigma, A8674 | 10 mM, DMSO, -20C | 0.5 μM |
| D-Glucose | Sigma, G8769 | 2.5M, 4C | 10 mM |
| 2-deoxyglucose | Sigma, D3179-1G | 500 mM, Glycolysis<br>Stress Test Media, -20C | 50 mM |
| Emetine | Sigma, E2375 | 100 mM, DMSO, -20C | 2 nM - 2500 nM |
| Puromycin | Sigma, P7255 | 10 mg/mL, DMSO, -20C | 1 mg/mL |
| Insulin | Sigma, 91077C | 1 mM, water, 4C | 1.6 nM - 1000 nM |
| Torin-2 | VWR, 103542-338 | 1 mM, DMSO, -20C | 1 nM - 250 nM |
| Rapamycin | VWR, 101762-276 | 50 mM, DMSO, -20C | 0.8 μM - 66 μM |

**Table 3.**
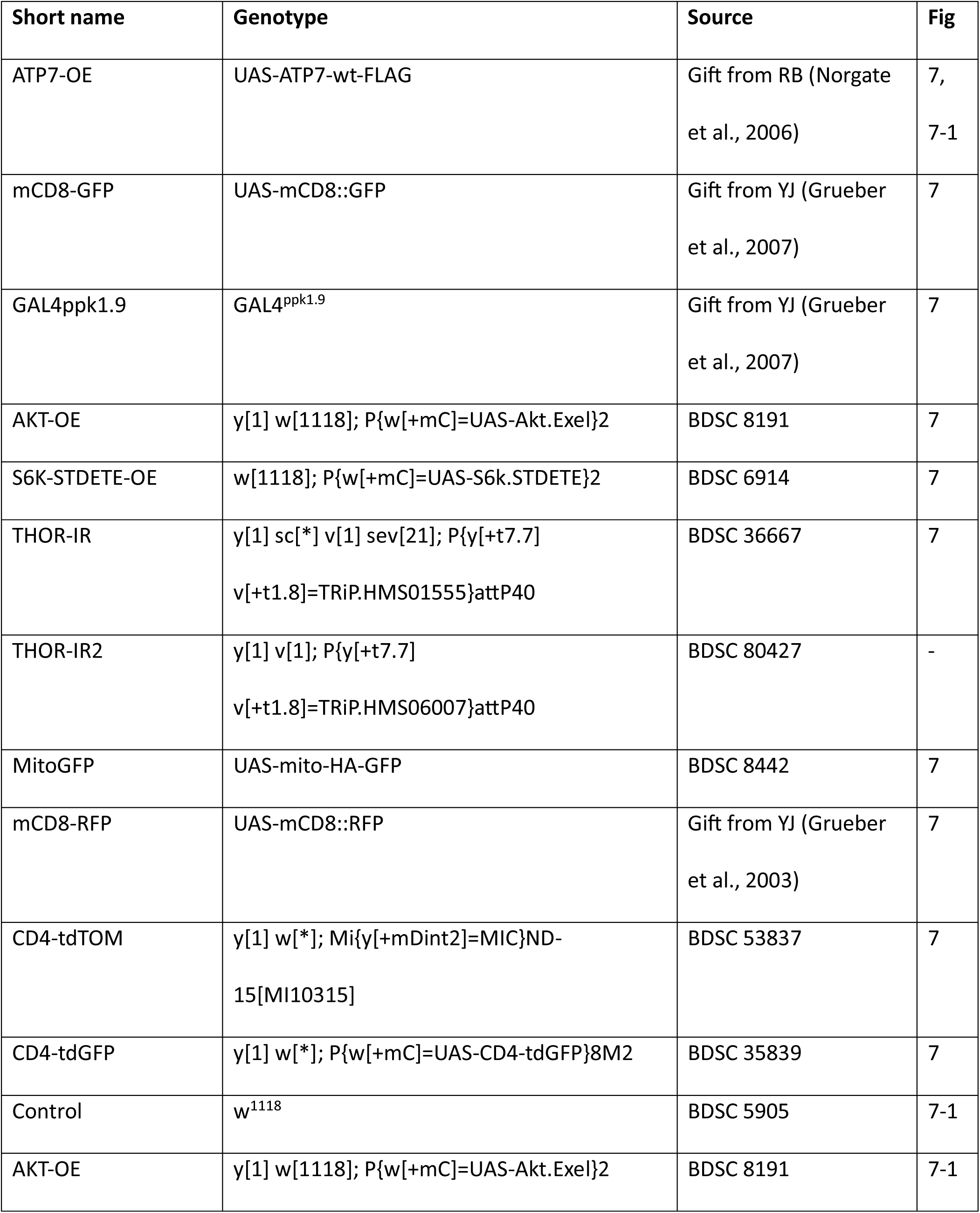
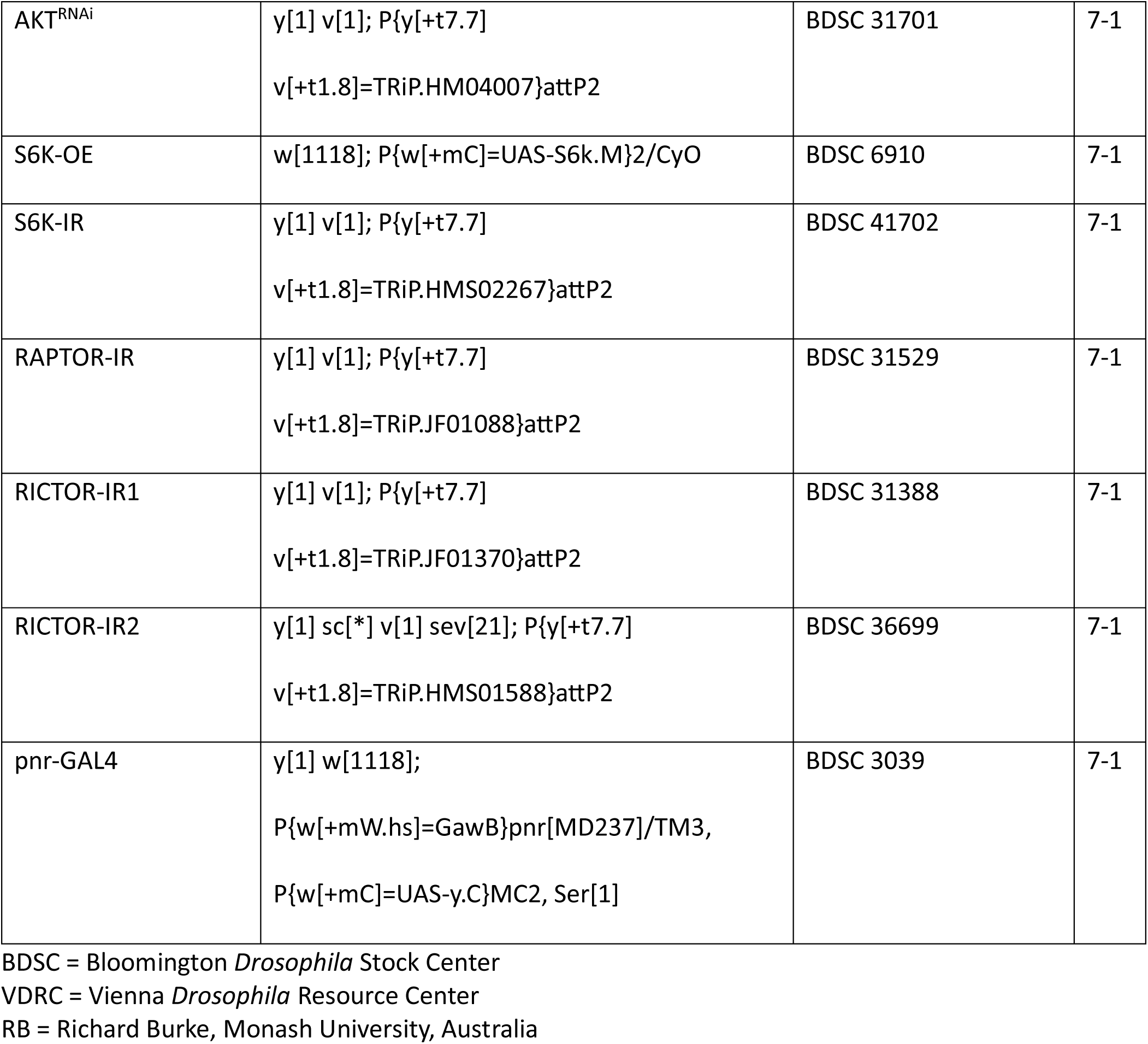
*Drosophila* Strains.

### Immunoblotting and puromycin pulse

Cells were grown in 12 or 24 well plates up to required confluency. Treatments are described in each figure. For puromycin pulse experiments, puromycin was added to the media 30 minutes before lysis to a final concentration of 1 mg/mL. The plates were placed on ice, and the cells were washed with cold phosphate-buffered saline (PBS) (Corning, 21-040-CV). Lysis buffer containing 150 mM NaCl, 10 mM HEPES, 1 mM ethylene glycol-bis(β-aminoethylether)- *N*,*N*,*N*′,*N*′-tetraacetic acid (EGTA), and 0.1 mM MgCl2, pH 7.4 (Buffer A), with 0.5% Triton X-100 (Sigma, T9284) and Complete anti-protease (Roche, 11245200) was added to each plate. For samples of interest for phosphorylated proteins, PhosSTOP phosphatase inhibitor (Roche, 04906837001) was also added to the lysis buffer. Cells were then scraped and placed in Eppendorf tubes on ice for 20 min and centrifuged at 16,100 × *g* for 10 min. The insoluble pellet was discarded, and the clarified supernatant was recovered. The Bradford Assay (Bio-Rad, 5000006) was used to determine protein concentration, and all lysates were flash frozen on dry ice and stored at -80C.

Cell lysates were reduced and denatured with Laemmli buffer (SDS and 2-mercaptoethanol) and heated for 5 min at 75C. Equivalent amounts of samples were loaded onto 4-20% Criterion gels (Bio-Rad, 5671094) for SDS-PAGE in running buffer (25 mM TRIS, 130 mM glycine, and 0.1% SDS) and transferred using the semidry transfer method with transfer buffer (48 mM TRIS, 39 mM glycine, 0.037% SDS, 20% methanol) to polyvinylidene difluoride (PVDF) membranes (Millipore, IPFL00010) unless otherwise specified in the figure legend that a nitrocellulose membrane was used (Sigma, 10600009). The membranes were incubated in TRIS-Buffered Saline (TBS: 1.36 M NaCl, 26.8 mM KCl, 247 mM TRIS) containing 5% nonfat milk and 0.05% Triton X-100 (TBST; blocking solution) for 30 min at room temperature (RT). The membrane was then rinsed thoroughly and incubated overnight with optimally diluted primary antibody in a buffer containing PBS with 3% bovine serum albumin (BSA) and 0.2% sodium azide. The next day, membranes were rinsed in TBST and treated with horseradish peroxidase-conjugated secondary antibodies against mouse or rabbit (see Table 1) diluted 1:5000 in the blocking solution for at least 30 min at RT. The membranes were washed in TBST at least three times and probed with Western Lightning Plus ECL reagent (PerkinElmer, NEL105001EA) and exposed to GE Healthcare Hyperfilm ECL (28906839). Additional staining was repeated as described above follows stripping of blots (200 mM glycine, 13.8 mM SDS, pH 2.5).

### Mitochondrial isolation and blue native gel electrophoresis

Starting material was two 150 mm dishes with cells at 80-90% confluency for each condition. The cells were released with trypsin and the pellet washed with PBS. Crude mitochondria were enriched according to Wieckowski et al. (2009). Briefly, cells were homogenized in isolation buffer (225 mM mannitol, 75 mM sucrose, 0.1 mM EGTA, and 30 mM Tris-HCl, pH 7.4) with 20 strokes in a Potter-Elvehjem homogenizer at 6000 rpm, 4°C. Unbroken cells and nuclei were collected by centrifugation at 600 × g for 5 min and mitochondria recovered from this supernatant by centrifugation at 7000 × g for 10 min. After one wash of this pellet, membranes were solubilized in 1.5 M aminocaproic acid, 50 mM Bis-Tris, pH 7.0, buffer with antiproteases and 4 g/g (detergent/protein) digitonin or DDM (n-dodecyl β-D-maltoside) to preserve or dissolve supercomplexes, respectively (Wittig et al., 2006; Timón-Gómez et al., 2020). Proteins were separated by blue native electrophoresis in 3-12% gradient gels (Novex, BN2011BX10; Díaz et al., 2009; Timón-Gómez et al., 2020) using 10 mg/ml ferritin (404 and 880 kDa, Sigma F4503) and BSA (66 and 132 kDa) as molecular weight standards.

### Cell survival and Synergy analysis

For all cell survival assays with a single drug, cells were counted using an automated Bio-Rad cell counter (Bio-Rad, TC20, 1450102) and plated in 96 well plates at 5,000-10,000 cells/well and allowed to sit overnight before drugs were added. Cells were treated with the drug concentrations indicated in each figure and summarized in Table 2 for 72 hours with the exception of emetine, for which double the number of cells were plated and cells were treated for 24 hours. Fresh media with 10% Alamar blue (Resazurin, R&D Systems #AR002) was added to each well and after 2 hours in the incubator absorbance was measured using a microplate reader (BioTek, Synergy HT; excitation at 530–570 nm and emission maximum at 580–590 nm) using a BioTek Synergy HT microplate reader with Gen5 software 3.11. For each experiment, percent survival was calculated by subtracting the background value of an empty well with only Alamar blue and normalizing to the untreated condition for each genotype. Individual data points represent the average survival of duplicate or triplicate treatments for each concentration.

For Synergy survival assays with two drugs, cells were plated in 96 well plates, treated with drugs, and incubated with Alamar blue as described above. Concentrations used are depicted in the corresponding figures and summarized in Table 2. For each experiment, percent survival was calculated by normalizing to the untreated condition for each genotype. For experiments with fetal bovine serum, values were normalized to the 10% serum condition, equivalent to the normal growth media. Individual data points represent the average survival of single replicates for each concentration. Synergy calculations were performed using the ZIP score with the SynergyFinder engine (https://synergyfinder.org/; Yadav et al., 2015; Zheng et al., 2022). For Torin 2-elesclomol experiments, the weighted ZIP score was used, a calculation which identifies synergy at lower drug doses with less toxicity by weighting the synergy distribution at each dose level using the proportion of responses from each drug in isolation (Ianevski et al., 2022).

### Seahorse metabolic oximetry

Extracellular flux analysis of the Mito Stress Test was performed on the Seahorse XFe96 Analyzer (Seahorse Bioscience) following manufacturer recommendations. SH-SY5Y cells were seeded at a density of 30,000 cells/well on Seahorse XF96 V3-PS Microplates (Agilent Technologies, 101085-004) after being trypsinized and counted (Bio-Rad TC20 automated Cell Counter) the day before the experiment. When appropriate, cells were treated with drugs in 10 cm plates before seeding in Seahorse XF96 microplates and treated overnight (see indicated times and concentrations in figure legends). XFe96 extracellular flux assay kit probes (Agilent Technologies, 102416-100) incubated with the included manufacturer calibration solution overnight at 37°C without CO_2_ injection. The following day, wells were washed twice in Seahorse Mito Stress Test Media. The Mito Stress Test Media consisted of Seahorse XF base media (Agilent Technologies, 102353-100) with the addition of 2 mM L-glutamine (HyClone, SH30034.01), 1 mM sodium pyruvate (Sigma, S8636), and 10 mM D-glucose (Sigma, G8769). After the washes, cells were incubated at 37°C without CO_2_ injection for 1 hour prior to the stress test. During this time, flux plate probes were loaded and calibrated. After calibration, the flux plate containing calibrant solution was exchanged for the Seahorse cell culture plate and equilibrated. Seahorse injection ports were filled with 10-fold concentrated solution of oligomycin A, FCCP, and rotenone mixed with antimycin A (see Table 2 for final testing conditions and catalog numbers). All Seahorse drugs were dissolved in DMSO and diluted in Seahorse Mito Stress Test Media for the Seahorse protocol. The flux analyzer protocol included three basal read cycles and three reads following injection of oligomycin A, FCCP, and rotenone plus antimycin A. Each read cycle included a 3-minute mix cycle followed by a 3-minute read cycle where oxygen consumption rate (OCR) and extracellular acidification rate (ECAR) were determined over time. In all experiments, OCR and ECAR readings in each well were normalized by protein concentration in the well. Cells were washed twice with phosphate buffered saline (Corning 21-040-CV) supplemented with 1 mM MgCl_2_ and 100 μM CaCl_2_ and lysed in Buffer A. Protein concentration was measured using the Pierce BCA Protein Assay Kit (Thermo Fisher Scientific, 23227) according to manufacturer protocol. The BCA assay absorbance was read by a BioTek Synergy HT microplate reader using Gen5 software. For data analysis of OCR and ECAR, the Seahorse Wave Software version 2.2.0.276 was used. Individual data points represent the average values of a minimum of three replicates. Non-mitochondrial respiration was determined as the lowest OCR following injection of rotenone plus antimycin A. Basal respiration was calculated from the OCR just before oligomycin injection minus the non-mitochondrial respiration. Non-mitochondrial respiration was determined as the lowest OCR following injection of rotenone plus antimycin A. ATP-dependent respiration was calculated as the difference in OCR just before oligomycin injection to the minimum OCR following oligomycin injection but before FCCP injection. Maximal respiration was calculated as the maximum OCR of the three readings following FCCP injection minus non-mitochondrial respiration.

Extracellular flux analysis of the Glycolysis Stress Test was performed as above with the following changes. The Glycolysis Stress Test Media contained only 2 mM L-glutamine. Seahorse injection ports were filled with 10-fold concentrated solution of D-glucose, oligomycin, and 2-Deoxy-D-glucose (2-DG) (see Table 2 for final testing conditions and catalog numbers). All drugs were diluted in Seahorse Glycolysis Stress Test Media for the Seahorse protocol. Glycolysis is calculated as the difference in ECAR between the maximum rate measurements before oligomycin injection and the last rate measurement before glucose injection. Glycolytic capacity was calculated as the difference in ECAR between the maximum rate measurement after oligomycin injection and the last rate measurement before glucose injection. Glycolytic reserve was calculated as the glycolytic capacity minus glycolysis. Non-glycolytic acidification was defined as the last rate measurement prior to glucose injection.

### Resipher

Poly-L-lysine coated Nunc 96 well (Thermo, 269787) or Falcon 96 well (Falcon, 353072) plates were seeded with 40,000 cells per well with 200 μL culture media (DMEM/10% FBS media). The following day, 150 μL of culture media was replaced with fresh, prewarmed, DMEM/10% FBS media. The Resipher sensing probe lid (Nunc Plates, NS32-N; Falcon Plates, NS32-101A) and Resipher system (Lucid Scientific, Atlanta, GA) were placed on one of the replicate plates and incubated in a humidified, 37C, 5% CO2 incubator while data was collected. For serum switch assays, at 48 hours after starting Resipher surveillance, 150 μL of warmed, un-supplemented DMEM media was replaced 3 times in 200 μL total volume to serially dilute assay to DMEM/0.16% FBS. For emetine assays, at approximately 48 hours after beginning Resipher surveillance, 150 μL of media was replaced with warmed DMEM/10% FBS media containing DMSO vehicle (VWR, WN182) or emetine (Sigma, E2375) to bring the final concentration in the wells to 3.2e-4% DMSO and 60 nM, 120 nM, or 240 nM emetine. After approximately 24 hours of DMEM/0.16%FBS or DMEM/10% FBS with vehicle/emetine treatment, rotenone (Sigma, R8875) and antimycin A (Sigma, A8674) were diluted to 10x in un-supplemented media and added to bring the well concentration to 1 mM rotenone and 1 mM Antimycin A. After about 24 hours of rotenone/antimycin treatment, the Resipher system assay was stopped. Cell counts were performed in parallel plates by measuring protein content at 24h, 48h (at the addition of emetine/DMSO/vehicle media), and 72h (at the addition of rotenone/antimycin A) and on the final Resipher plate after respiration was stably down following the addition of rotenone and antimycin A. To determine protein concentrations, wells were washed three times with phosphate buffered saline (Corning 21-040-CV) supplemented with 1 mM MgCl2 and 100 μM CaCl2, and cells were lysed in Buffer A with Complete antiprotease. Protein concentration was measured using the Pierce BCA Protein Assay Kit (Thermo Fisher Scientific, 23227) according to manufacturer protocol. Experiments were repeated in quadruplicate. Doubling time (*T_d_*) was estimated using the equation N(t)=N(0)2^t/*Td*^ where N(0) is the protein at either 24 or 48h and N(t) is the protein 24h later. Accumulated cell counts were estimated by determining areas under the curve using Prism Version 10.2.2.

### Total RNA extraction and NanoString mRNA Quantification

Cells were grown on 10 cm plates, and total RNA was extracted using the TRIzol reagent (Invitrogen, 15596026). When applicable, cells were treated with 200 μM BCS for 3 days. For preparation of samples, all cells were washed twice in ice-cold PBS containing 0.1 mM CaCl2 and 1.0 mM MgCl2. 1 ml of TRIzol (Invitrogen, 15596026) was added to the samples and the TRIzol mixture was flash frozen and stored at -80C for a few weeks until RNA Extraction and NanoString processing was completed by the Emory Integrated Genomics Core. The Core assessed RNA quality before proceeding with the NanoString protocol. The NanoString Neuropathology gene panel kit (XT-CSO-HNROP1-12) or Metabolic Pathways Panel (XT-CSO-HMP1-12) was used for mRNA quantification. mRNA counts were normalized to either the housekeeping genes AARS or TBP, respectively, using NanoString nSolver software. Normalized data were further processed and visualized by Qlucore.

### ICP mass spectrometry

Procedures were performed as described previously (Lane et al., 2022). Briefly, cells were plated on 10 or 15 cm dishes. After reaching desired confluency, cells were treated with 1 nM elesclomol for 24 hours. On the day of sample collection, the plates were washed three times with PBS, detached with trypsin, and neutralized with media and pelleted at 800 × g for 5 min at 4C. The cell pellet was resuspended with ice-cold PBS, aliquoted into 3-5 tubes, centrifuged at 16,100 × g for 10 min. The supernatant was aspirated, and the residual pellet was immediately frozen on dry ice and stored at -80C. Mitochondria were isolated as described above. Tissue samples were collected following euthanasia, weighed, and immediately flash frozen on dry ice. Cell or mitochondrial pellets were digested by adding 50 µL of 70% trace metal basis grade nitric acid (Millipore Sigma, 225711) followed by heating at 95C for 10 min. Tissue was digested by adding 70% nitric acid (50-75% w/v, i.e. 50 mg of tissue was digested in 100 µL of acid) and heated at 95C for 20 min, followed by the addition of an equal volume of 32% trace metal basis grade hydrogen peroxide (Millipore Sigma, 95321) and heated at 65C for 15 minutes. After cooling, 20 µL of each sample was diluted to 800 µL to a final concentration of 2% nitric acid using either 2% nitric acid or 2% nitric acid with 0.5% hydrochloric acid (VWR, RC3720-16) (vol/vol). Metal levels were quantified using a triple quad ICP-MS instrument (Thermo Fisher, iCAP-TQ) operating in oxygen mode under standard conditions (RF power 1550 W, sample depth 5.0 mm, nebulizer flow 1.12L/min, spray chamber 3C, extraction lens 1,2 -195, –-15 V). Oxygen was used as a reaction gas (0.3 mL/min) to remove polyatomic interferences or mass shift target elements (analytes measured; 32S.16O, 63Cu, 66Zn). External calibration curves were generated using a multielemental standard (ICP-MSCAL2-1, AccuStandard, USA) and ranged from 0.5 to 1000 µg/L for each element. Scandium (10 µg/L) was used as internal standards and diluted into the sample in-line. Samples were introduced into the ICP-MS using the 2DX PrepFAST M5 autosampler (Elemental Scientific) equipped with a 250 µL loop and using the 0.25 mL precision method provided by the manufacturer. Serumnorm (Sero, Norway) was used as a standard reference material, and values for elements of interest were within 20% of the accepted value. Quantitative data analysis was conducted with Qtegra software, and values were exported to Excel for further statistical analysis.

### Preparation of brain tissue for proteomics, immunoblots, or Luminex analysis

Brain samples were collected after euthanasia and immediately flash frozen in liquid nitrogen and stored at -80C. Brains were lysed in 8M urea in 100 μM potassium phosphate buffer (pH 8.0; 47.6 mL 1M K_2_HPO_4_ + 4.8 mL 1M KH_2_PO_4_, bring to 525 mL total volume) with Complete anti-protease and PhosSTOP phosphatase inhibitor and homogenized by sonication (Fisher Scientific, Sonic Dismembrator Model 100). After incubation on ice for 30 minutes, samples were spun at 7000 × g for 10 minutes at 4C, and the supernatant was transferred to a new tube. Protein concentration was measured in triplicate using the Pierce BCA Protein Assay Kit (Thermo Fisher Scientific, 23227) according to manufacturer protocol. Samples were stored at -80C until use.

### TMT mass spectrometry for proteomics

Cells were grown in standard media as described above (Fig. 2C, TMT1; DMEM (Corning, 10-013), 10% FBS, 25 mM D-glucose, 4 mM L-glutamine, 1 mM sodium pyruvate) or media with dialyzed FBS supplemented with D-glucose, sodium pyruvate, and L-glutamine (Fig. 2C, TMT2; DMEM (Thermo Fisher, A14430-01), 10% dialyzed FBS (Thermo Fisher, 26400-044), 10 mM D-glucose, 2 mM L-glutamine, 1 mM pyruvate). There was no difference in total cellular copper as measured by ICP-MS (not shown; manuscript in preparation containing the complete TMT2 dataset). Cells were detached with PBS-EDTA (ethylenediaminetetraacetic acid, 10 mM) and pelleted as described above for ICP-MS. The supernatant was aspirated, and the pellet was immediately frozen on dry ice and stored at -80C.

TMT labeling was performed according to the manufacturer’s protocol and as described previously (Ping et al., 2018; Wynne et al., 2023). Briefly, samples were lysed in urea buffer (8 M urea, 100 mM NaH_2_PO_4_, pH 8.5; 1X HALT protease and phosphatase inhibitor cocktail, Pierce) and underwent probe sonication. Protein concentration was measured by BCA assay, and aliquots were prepared with 100 μg lysate in the same total volume of lysis buffer. Aliquots were diluted with 50 mM HEPES (pH 8.5) and sequentially treated with 1 mM DTT (1,4-dithiothreitol) and 5 mM IAA (iodoacetamide) for alkylation at room temperature, followed by overnight digestion with lysyl endopeptidase (Wako) at a 1:50 (w/w) enzyme to protein ratio, dilution to a urea concentration of 1 M with 50 mM triethylammonium bicarbonate (TEAB), and overnight digestion with trypsin (Promega) at a 1:50 (w/w) enzyme to protein ratio. Peptides were desalted with a Sep-Pak C18 column (Waters) and aliquots of each sample (equivalent to 20 μg protein per sample) were combined to obtain a global internal standard (GIS) for TMT labeling. All 20 samples were then dried under vacuum (16 individual and 4 GIS), resuspended in 100 mm TEAB buffer followed by the addition of anhydrous acetonitrile, and solutions were transferred to their respective channel tubes. After 1 hr, the reaction was quenched with 5% hydroxylamine and all samples were combined and dried. Samples were resuspended in 90% acetonitrile and 0.01% acetic acid and loaded onto an offline electrostatic repulsion–hydrophilic interaction chromatography fractionation HPLC system. 96 fractions were collected, combined into 24 fractions, dried down, and resuspended in peptide loading buffer (0.1% formic acid, 0.03% trifluoroacetic acid, 1% acetonitrile). Peptide mixtures (2 μL) were separated on a self-packed C18 fused silica column by an Easy-nLC 1200 and monitored on a Fusion Lumos mass spectrometer (Thermo Fisher Scientific). MS/MS spectra were searched against the Uniprot human database (downloaded on 04/2015) with Proteome Discoverer 2.1.1.21 (Thermo Fisher Scientific). Variable and static modifications included methionine oxidation, asparagine, glutamine deamidation, protein N-terminal acetylation, cysteine carbamidomethyl, peptide N-terminus TMT, and lysine TMT. Percolator was used to filter MS/MS spectra matches false discovery rate of <1%. Abundance calculations used only razor and unique peptides, and the ratios of sample over the GIS of normalized channel abundances were used for comparison across all samples. Data files will be uploaded to ProteomeExchange.

### Metabolite quantification by LC mass spectrometry

Cells were seeded in duplicate 6 wells per replicate at 100,000 or 190,000 cells per well for wild-type and CTR1 KO cells, respectively, and treated with 1 nM elesclomol for 2 days. Media was replaced with pyruvate-free media (Thermo Fisher 11966-025, supplemented with 25 mM glucose) with dialyzed FBS (Thermo Fisher 26400044) and 1 nM elesclomol and 5 μM CuCl_2_ for 24 hours. (Media prepared with dialyzed FBS must be supplemented with copper as dialysis removes copper from the serum.) After a total of 72h elesclomol treatment, cells were washed with ice-cold PBS supplemented with 1 mM MgCl_2_ and 100 μM CaCl_2_ and extracted with 400 μL of lysis buffer (0.1M formic acid at 4:4:2 dilution (MeOH: ACN: Water) containing 25 µM ^13^C_3_ alanine as an internal standard. After 2 min on ice, 35 μL of 15% NH_4_HCO_3_ was added, mixed well by gently swirling the plate, and incubated for another 20 min on ice. Cell lysates were then transferred into pre-chilled 1.5 ml centrifuge tubes, vortexed briefly, and spun at 21,300xg for 30 min at 4°C. 360 μL of supernatant was then transferred into pre-chilled 1.5 ml centrifuge tubes and dried down using a Savant Speedvac Plus vacuum concentrator. Samples were resuspended in 120 μL of 60:40 (ACN: Water), sonicated for 5 minutes at 4°C, and centrifuged at 21,300xg for 20 min at 4°C. 100 μL of supernatant was transferred into a pre-chilled LC-MS vial. 5 μL of this sample was injected into a HILIC-Z column (Agilent Technologies) on an Agilent 6546 QTOF mass spectrometer coupled with an Agilent 1290 Infinity II UHPLC system (Agilent Technologies). The column temperature was maintained at 15 °C and the autosampler was at 4 °C. Mobile phase A: 20 mM Ammonium Acetate, pH = 9.3 with 5 μM Medronic acid, and mobile phase B: acetonitrile. The gradient run at a flow rate of 0.4 ml/min was: 0min: 90% B, 1min 90% B, 8 min: 78% B, 12 min: 60% B, 15 min: 10% B, 18 min: 10 %B, 19-23 min: 90% B. The MS data were collected in the negative mode within an m/z = 20-1100 at 1 spectrum/sec, Gas temperature: 225 °C, Drying Gas: 9 l/min, Nebulizer: 10 psi, Sheath gas temp: 375 °C, Sheath Gas flow: 12 l/min, VCap: 3000V, Nozzle voltage 500 V, Fragmentor: 100V, and Skimmer: 45V. Data were analyzed using Masshunter Qualitative Analysis 10 and Masshunter Quantitative Analysis 11 (Agilent Technologies). Metabolite levels from different treatments were normalized to cell numbers.

### Insulin receptor phosphorylation quantification

The cerebellum was isolated from mice at postnatal day 10 and flash frozen in liquid nitrogen and stored at -80C. Tissue was dissolved in 300 μL 8M urea in 100 mM PO_4_ containing protease and phosphatase inhibitors, sonicated 5-10 times in 1 second bursts, and incubated on ice for 30 minutes with periodic vortexing. Samples were spun at 13,500 RCF for 10 min at 4C and the supernatant was transferred to a new tube. Protein concentration was measured in triplicate by BCA as described above. Mouse brain lysates were stored at -80°C, then thawed on ice and normalized to 1µg of total protein in Milliplex Assay buffer prior to the start of the assay protocol. We measured Tyr1135/Tyr1136 phosphosites in IGF1R with the Milliplex® MAP kit (48-611MAG) read out on a MAGPIX Luminex instrument (Luminex, Austin, TX, USA).

### Digital spatial profiling of mouse brain tissue

Formalin-fixed paraffin-embedded mouse brain tissue was profiled using GeoMx^®^ DSP (Merritt et al., 2020). Brains were isolated at P10 after euthanasia and immediately fixed overnight at room temperature in 10% neutral buffered formalin (Thermo Fisher 28906, diluted to 10% NBF with 0.9% sterile saline solution) at a ratio of 20:1 fixative to sample. Thin (5 µm) sagittal tissue sections were prepared on positively charged slides by the Emory University Cancer Tissue and Pathology shared resource according to manufacturer’s recommendations for Semi-Automated RNA Slide Preparation Protocol (FFPE) (manual no. MAN-10151-04). Sections were air-dried at room temperature overnight and shipped to NanoString at room temperature.

Following an overnight bake at 65°C, slides underwent deparaffinization, rehydration, heat-induced epitope retrieval (for 20 minutes at 100°C with Bond Epitope Retrieval 2 Solution), and enzymatic digestion (0.1 μg/mL proteinase K for 15 minutes at 37°C). Tissues were then incubated with 10% neutral buffered formalin for 5 minutes and 5 minutes with NBF Stop buffer. All steps following overnight baking were carried out on a Leica BOND-RX. Slides were then removed from the Leica BOND-RX and *in situ* hybridization with GeoMx^®^ Mouse Whole Transcriptome Atlas (mWTA) probes was carried out overnight in a humidified hybridization chamber kept at 37°C. Following rounds of stringent washing with a 1:1 volumetric mixture of 4X SSC and 100% formamide to remove off-target probes, the tissue was blocked with Buffer W blocking solution (NanoString Technologies) then incubated with a mouse anti-GFAP antibody (see Table 2) and a stain for the nuclear marker Syto83.

Tissue morphology was visualized using fluorescent antibodies and Syto83 on the GeoMx^®^ DSP instrument. Regions of interest (ROIs) were selected from the cerebellum of the samples using the polygon tool. Either a GFAP+ or GFAP-mask was generated within each ROI using the fluorescence signal associated with the GFAP antibody, and UV light was utilized to release and collect oligonucleotides from each ROI. Areas of UV light irradiation within an ROI are referred to as areas of illumination (AOI). During PCR, Illumina i5 and i7 dual-indexing primers were added to each photocleaved oligonucleotide allowing for unique indexing of each AOI. Library concentration was measured using a Qubit fluorometer (Thermo Fisher Scientific), and quality was assessed using a Bioanalyzer (Agilent). The Illumina Novaseq 6000 was used for sequencing, and the resulting FASTQ files were then processed by the NanoString DND pipeline to generate count data for each target probe in every AOI.

### Quality control and preprocessing of GeoMx transcript data

The GeoMx DSP Analysis Suite was utilized for conducting both quality control (QC) and data exploration. First, each AOI was QC checked to ensure contamination was avoided during PCR and library preparation and that sequencing was sufficient. All AOIs passed QC so none were removed from downstream analysis. Next, any global outliers in the target list were identified and removed from the panel. Finally, the dataset was normalized using third quantile (Q3) normalization. The QC checked, Q3 normalized dataset was used for downstream analysis.

### Drosophila husbandry and genotypes

Fly strains used are listed in Table 3. All fly strains were reared at 25C on standard molasses media (Genessee Scientific) on a 12hr:12hr light:dark cycle. All fly strains were isogenized and bred into the same genetic background. For epidermis experiments (Extended Data Fig. 7-1), flies were aged 4-7 days and imaged using an Olympus SZ-61 stereomicroscope and an Olympus DP23 color camera. Images were then stacked using Zyrene stacker. Dendritic characterization experiments were performed in both male and female flies as described previously (Hartwig et al., 2020). Briefly, virgin females from mCD8-GFP were outcrossed to male flies from individual transgenic fly lines (AKT-OE, S6K-STDETE-OE, THOR-IR, and THOR-IR2) to first determine their effects on dendritic morphology. Of the two Thor RNAi lines tested, THOR-IR was the most phenotypic and all further experiments were performed using this line. The effect of ATP7-OE was determined using the ATP7-OE;mCD8-GFP transgenic fly line. Rescue analyses were performed by outcrossing ATP7-OE;mCD8-GFP virgin female flies to male flies from the fly lines previously mentioned. For both mCD8-GFP and ATP7-OE;mCD8-GFP, CD4-tdGFP was used as control. For super-resolution imaging, UAS-mito-HA-GFP;GAL4ppk1.9,UAS-mCD8::GFP virgin female flies were outcrossed to male flies from UAS-ATP7, UAS-ATP7;UAS-S6k.STDETE, or UAS-CD4-tdTOM (control).

### Drosophila dendritic imaging and analysis

Live confocal imaging was done as previously described (Hartwig et al., 2020). Neurons were imaged from wandering third-instar larvae on a Zeiss LSM780 microscope. Individual animals were placed on a microscopic slide and anesthetized by immersion in 1:5 (v/v) diethyl ether to halocarbon oil solution and covered with 22 x 50 mm coverslip. Images were acquired as z-stacks using a 20x dry objective (NA 0.8) and step size 2μm. Maximum intensity projections of the images were acquired, and the images were exported as .jpeg files using the Zen Blue software. Images were then stitched using Adobe Photoshop and manually curated using the Flyboys software to remove background noise (Das et al., 2017). The images were skeletonized and processed on ImageJ (Schneider et al., 2012; Arshadi et al., 2021) using a custom macro (https://github.com/CoxLabGSU/Drosophila_Sensory_Neuron_Quantification) to get quantitative metrics such as total dendritic length, branches, maximum intersection (Sholl), radius of maximum intersection (Sholl), and dendritic field coverage (convex hull). For proximal-distal analysis, a circular region of 400 pixels diameter from the soma was selected and considered as proximal to the soma. The region beyond 400 pixels was considered as distal. These regions were then analyzed separately. Average branch length was calculated by dividing the total dendritic length by the branches. The data obtained were then compiled using R6 (R Core Team, 2021) and exported to Excel (Microsoft).

Super-resolution imaging was performed on a Zeiss LSM980 Airyscan 2 confocal microscope on live animals. Animals were prepared as described above, and images were acquired as z-stacks using a 63x oil objective on Airyscan mode (SuperResolution:9.0 (3d, Auto)). Maximum intensity projections of the images were created using the Zen Blue software and exported as .TIFF files.

### Data availability

The mass spectrometry proteomics data will be deposited to the ProteomeXchange Consortium via the PRIDE (Deutsch et al., 2020) partner repository.

### Bioinformatic analyses and statistical analyses

Transcriptomics and proteomics data were processed with Qlucore Omics Explorer Version 3.6(33). Data were normalized to a variance of 1 and an average of 0 for statistical analysis and thresholding. Permutation statistical analyses were performed with the engine https://www.estimationstats.com/#/ with a two-sided permutation t-test using 5000 reshuffles (Ho et al., 2019). ANOVA and paired analyses were conducted with Prism Version 10.2.2 (341). Gene ontology studies were performed with Metascape and ENRICHR (Zhou et al., 2019).

## Author Contributions

Conceptualization, VF, AL, and EW; Methodology and Validation, AL, SB, SZ, AR, DD, LW, AP, AV, DC, BR, EW; Formal Analysis, VF, AL, and MM; Investigation, VF, AL, NS, SB, SZ, AR, AG, KS, DD, MM, WL, AB, FRM, TT, AP, LC, EW; Resources, VF, AL, SB, DC, MP, AV,; Writing – Original Draft, VF, AL, SB, MM, FRM, AV; Writing – Review & Editing, VF, AL, SB, SZ, AG, KS, MM, FRM, MP, LW, AP, AV, DC, EW; Visualization, VF, AL and MM; Supervision, VF, EW, LW, AP, AV, DC, BR; Funding Acquisition, VF, AL, KS, DC, AP, EW.

## Acknowledgements

This work was supported by NIH grants 1RF1AG060285 to VF, 1F31NS127419 to ARL, 4K00NS108539 to KSS, T34GM131939 to TT, R01NS086082 to DNC, 1R01NS133344-01A1 to AP, and R01ES034796 to EW and a Burroughs Wellcome 2021 Postdoctoral Grant to KSS. ARL is supported by an ARCS Foundation Award, Robert W. Woodruff Fellowship, John B. Lyon Memorial Scholarship Award, and NanoString Young Investigator Brain Tank award. This study was supported in part by the Emory Integrated Genomics and Proteomics Cores, which are subsidized by the Emory University School of Medicine, and the Cancer Tissue and Pathology shared resource of Winship Cancer Institute of Emory University and NIH/NCI under award number P30CA138292. ARL is grateful for personal support from LTR and D. VF is grateful for mitochondria provided by Maria Olga Gonzalez.

## Extended Data

**Figure 1-1.**
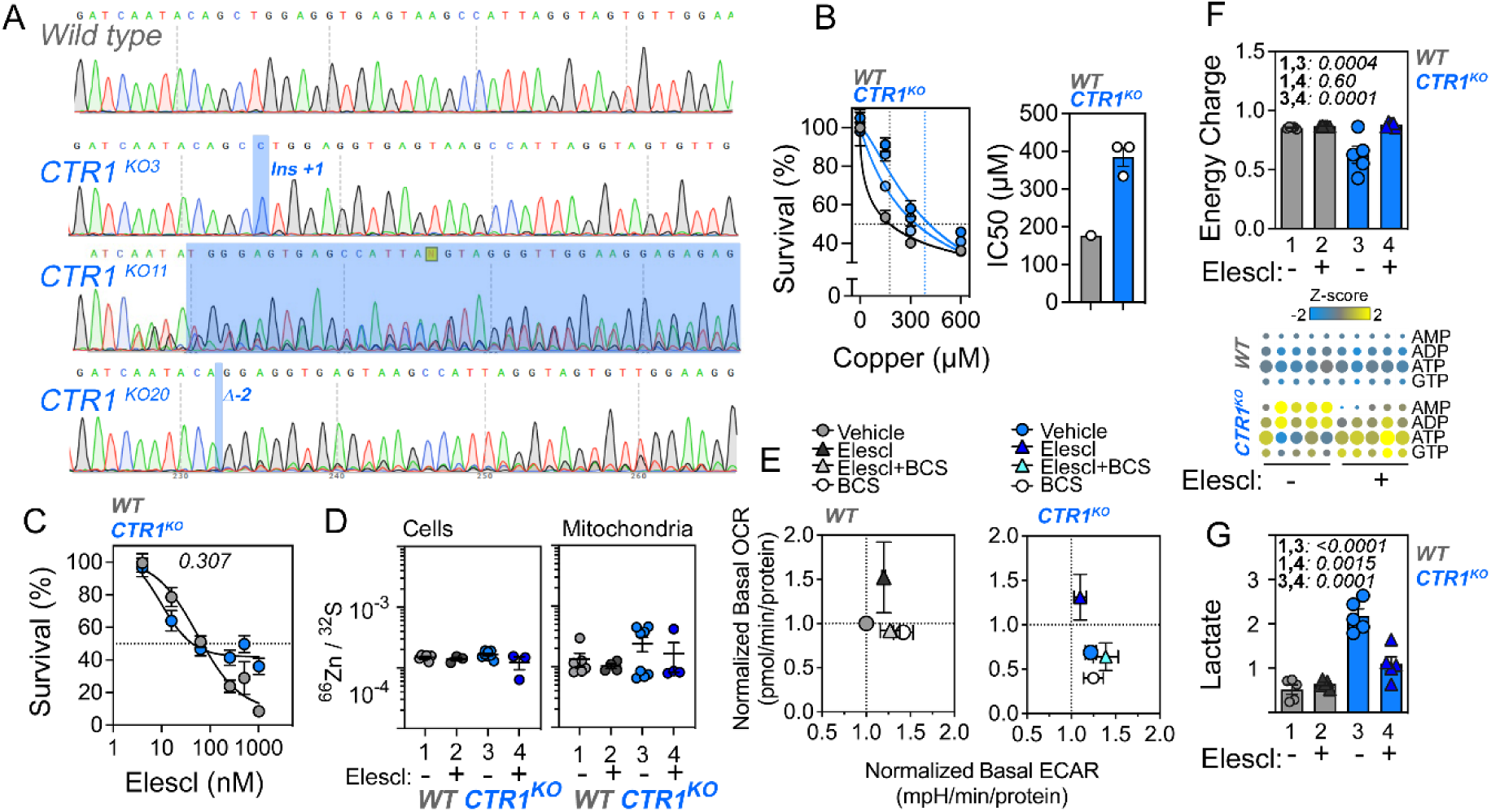
Generation and Characterization of CTR1 (*SLC31A1*) Null Mutant Neuroblastoma Cells. **A.** DNA sequence chromatograms of one wild type and three *SLC31A1* CRISPR-edited SH-SY5Y clonal lines. Blue boxes mark the mutated sequence. **B.** Cell survival analysis of *SLC31A1* mutants with increasing concentrations of copper. Blue shaded symbols represent three different clones (n=3 replicates for 1 or 3 clones for wild type or *SLC31A1* mutants respectively). IC50 was calculated by fitting data to a non-linear model using Prism. **C.** Cell survival analysis of wild type and CTR1 mutants with increasing concentrations of elesclomol (n = 11 and 14, p value Sum-of-squares F test analysis). **D.** 64Zn quantification in whole cells or mitochondria in CTR1 KO cells treated with vehicle or 1 nM elesclomol, normalized to 32S. There are no significant differences by treatment or genotype as assessed by One-Way ANOVA, followed by Benjamini, Krieger, and Yekutieli multiple comparisons correction. **E.** Metabolic map of basal OCR and ECAR from the Mito Stress Test of wild-type and CTR1 KO cells (clone KO20) normalized to untreated wild-type cells treated with vehicle, elesclomol, or BCS as described in Fig. 1. **F-G**. Mass spectrometry quantification of metabolites, analyzed by One-Way ANOVA followed by Benjamini, Krieger, and Yekutieli multiple comparisons correction. CTR1 clone KO20 was used. **G**. Quantification of energy charge, calculated by dividing the amount of [ATP] + 0.5[ADP] by the total levels of [ATP]+[ADP]+[AMP] (Atkinson and Walton, 1967). Heat maps show the levels of the adenylate nucleotides used for the calculation and GTP for comparison. **G.** Quantification of lactate levels. All data are presented as average ± SEM.

**Figure 2-1.**
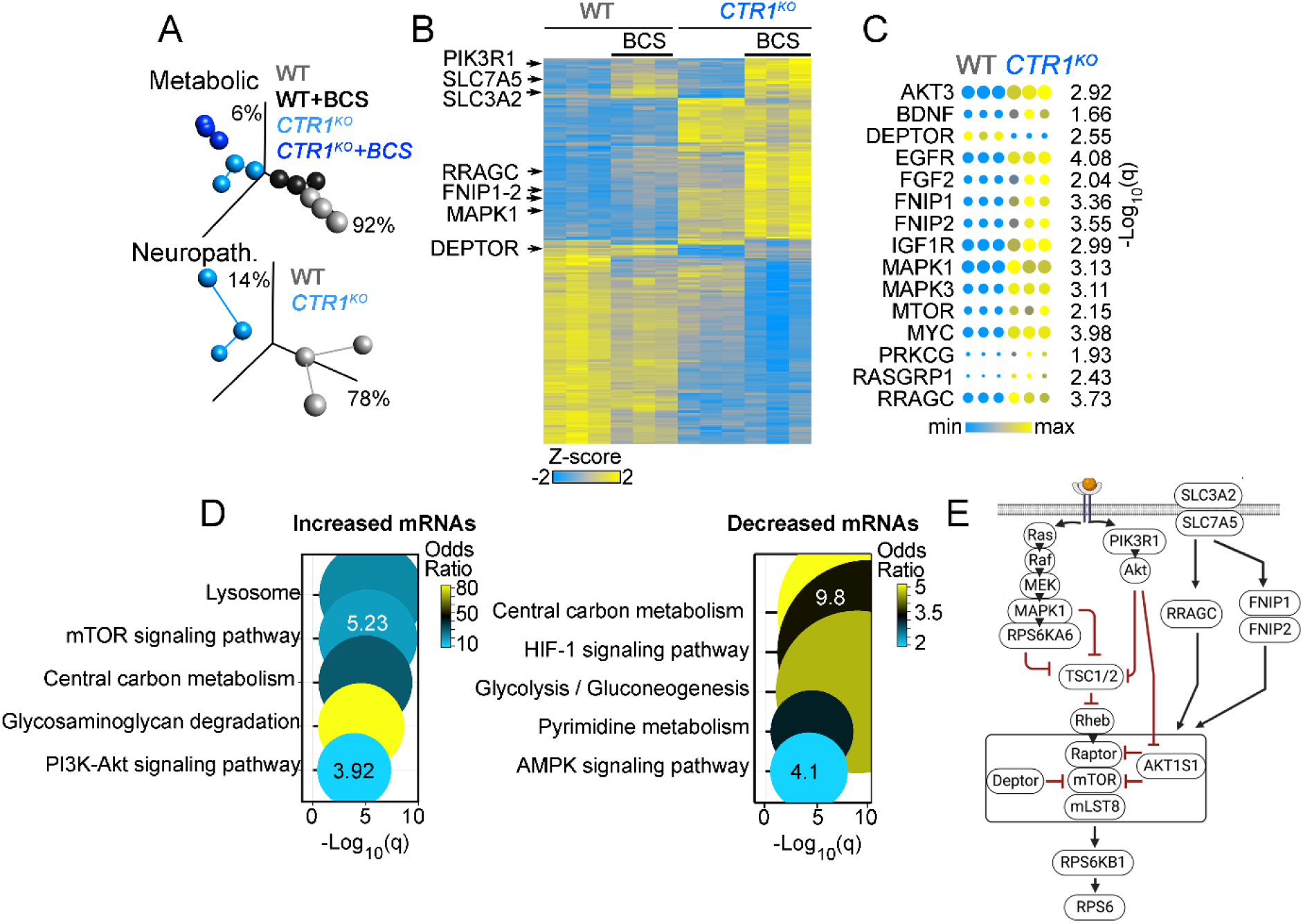
Transcript changes in CTR1 mutants reflect increased activation of the mTOR and PI3K-Akt signaling pathways. **A.** Principal component analysis of the Metabolic and Neuropathology annotated transcriptomes from wild type (gray) or CTR1 KO (blue symbols; KO20) clonal lines in the absence or presence of BCS (200 μM, 72 hours). **B.** Hierarchical clustering of the metabolism annotated transcriptome differentially expressed at a significancy of q<0.05 (analyzed by one-way ANOVA followed by Benjamini-Hochberg FDR correction). **C.** mTOR pathway annotated transcript levels, expressed as relative level. Each of these transcripts changed >1.5-fold as compared to wild type cells (q<0.05). **D.** Gene ontology analysis of metabolism annotated transcripts with increased or decreased expression in CTR1 mutants. KEGG database was queried with the ENRICHR engine. Fisher exact test followed by Benjamini-Hochberg correction. **E.** mTOR signaling pathway diagram modified from KEGG map04150. See Extended Data File 1 for raw data.

**Figure 4-1.**
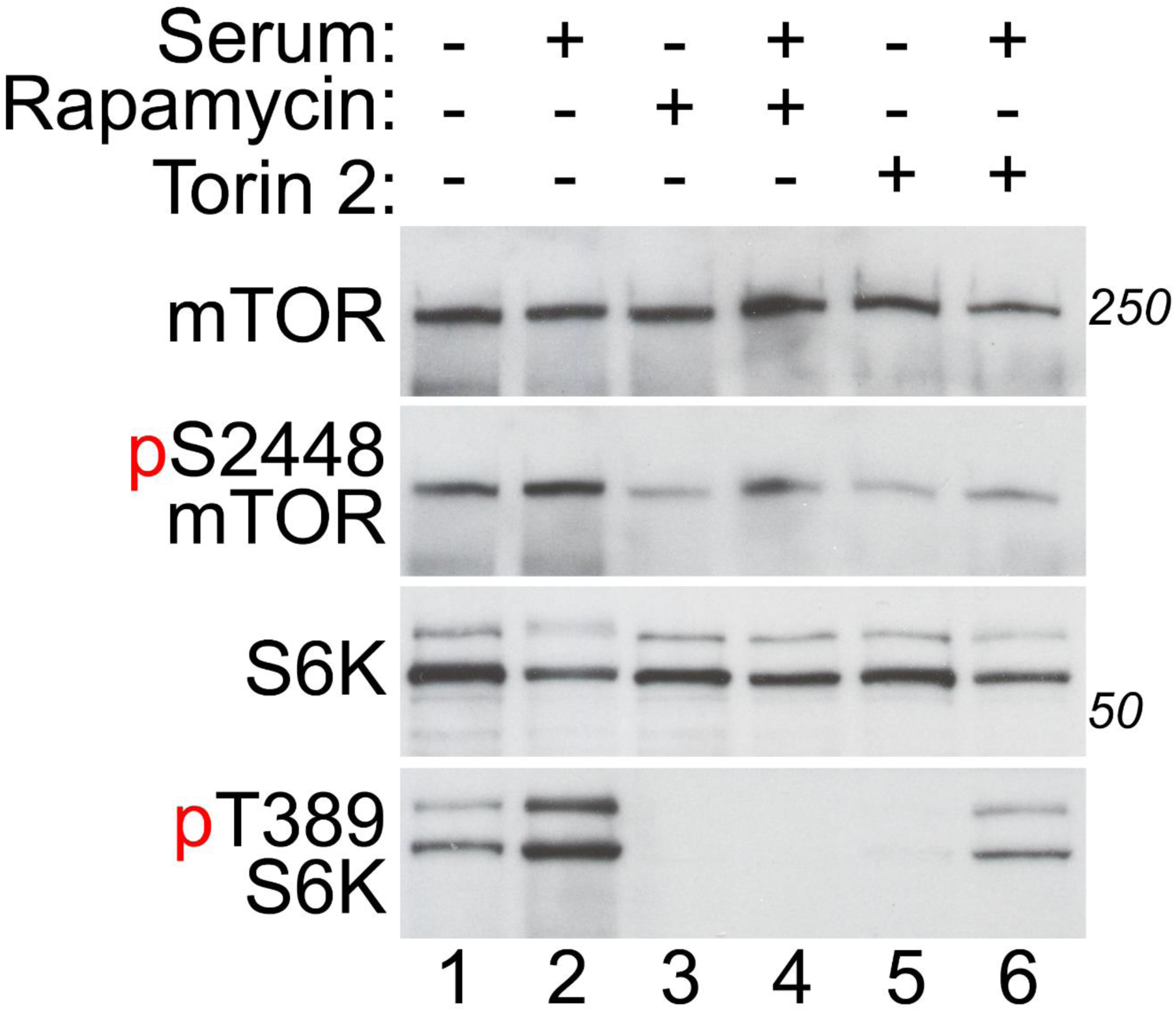
mTOR inhibitors reduce phosphorylation of mTOR and S6K. Immunoblot of total and phosphorylated mTOR and S6K after overnight serum depletion in 0.2% serum, 30 minute pretreatment with rapamycin (270 nM) or Torin-2 (1 nM) in existing 0.2% serum media, followed by a switch to fresh media with 0.2% or 20% serum, rapamycin, and/or Torin-2 for 30 minutes.

**Figure 4-2.**
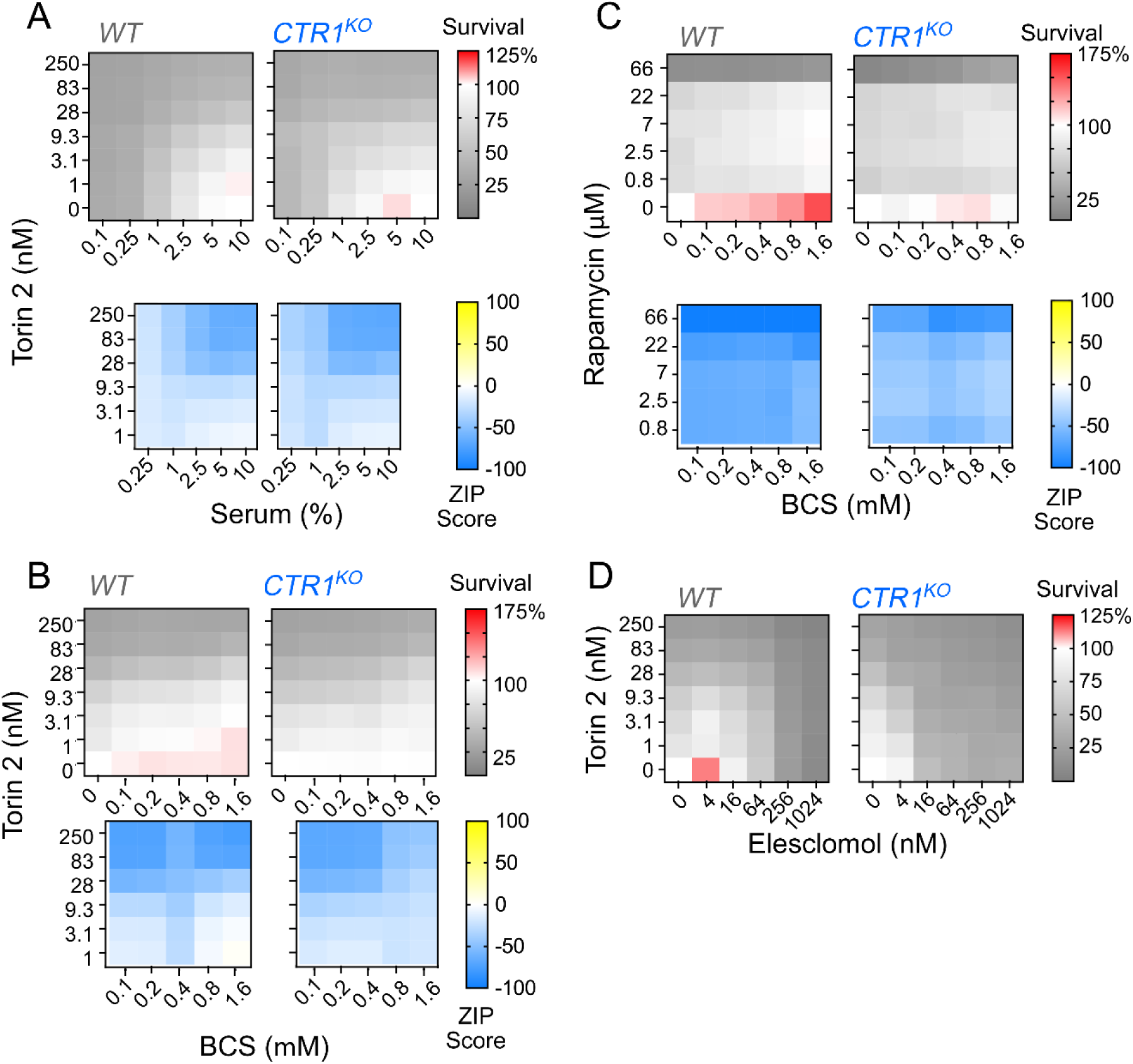
Synergy analysis of CTR1 mutant cells. **A-D**. Quantification of survival and drug synergy in wild-type and CTR1 mutant cells treated with Torin-2 and serum (**A**), Torin-2 and BCS (**B**), rapamycin and BCS (**C**), or Torin-2 and elesclomol (**D**). Gray panels represent cell survival normalized to untreated cells, as described in Figure 4. Blue panels represent the corresponding interaction synergy map calculated using the Zero Interaction Potency (ZIP) score for cell survival (Yadav et al., 2015) (refer to Figure 4).

**Figure 5-1.**
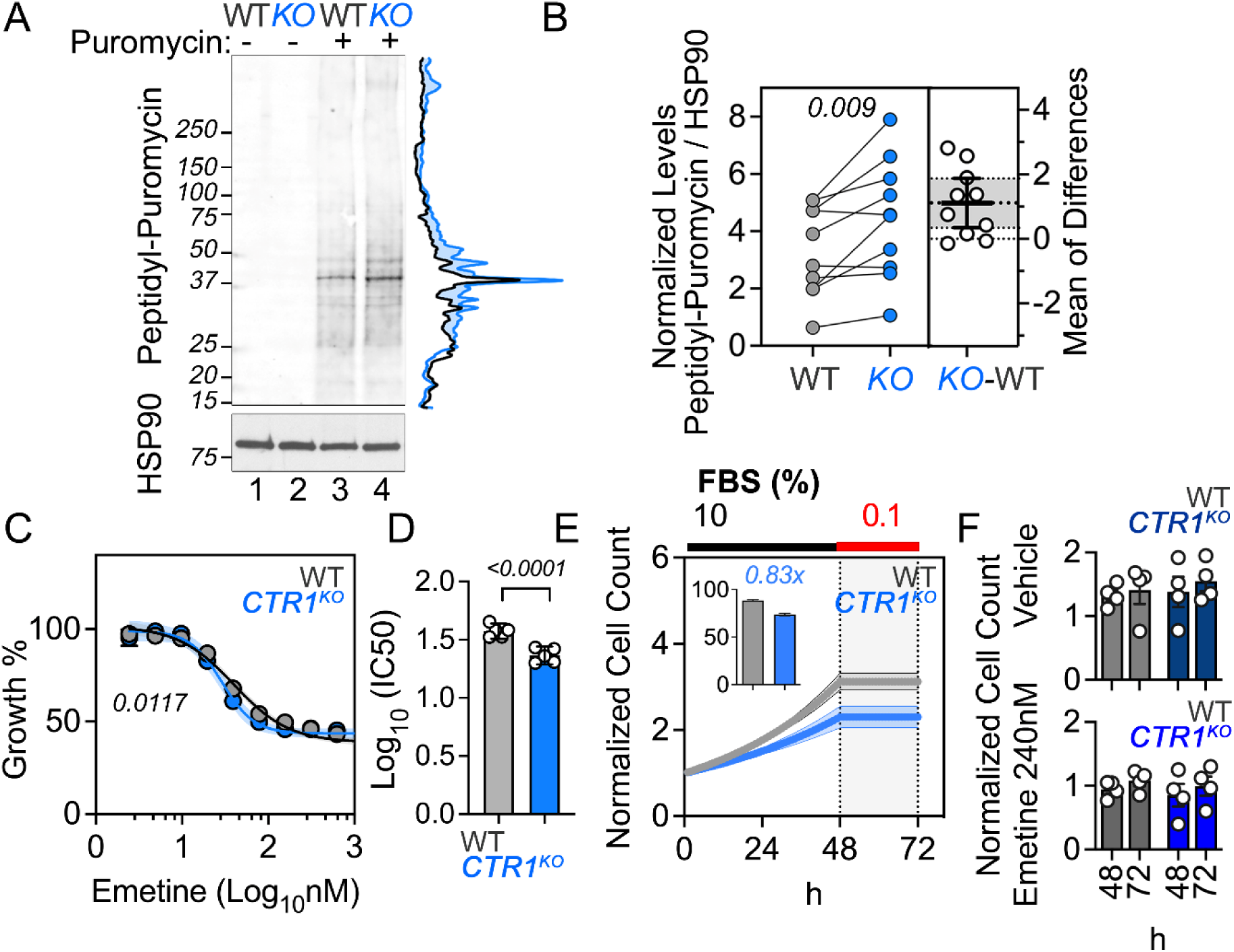
Cell survival under conditions inhibiting protein synthesis. **A-B.** Immunoblot for puromycin using a nitrocellulose membrane in wild-type and CTR1 mutant cells treated with vehicle (lanes 1 and 2) or puromycin for 30 min (lanes 3 and 4). Line trace represents the corresponding quantification of signal intensity. **B.** Quantification of the puromycin signal between 250 and 15 kDa normalized to HSP90. Paired two-tailed t test (pairing r=0.86, p=0.0007). **C-D.** Cell growth analysis of wild type and CTR1 mutants with increasing concentrations of emetine for 24 hours (**D**, Average ± SEM, n = 6, p value calculated with Sum-of-squares F test analysis) and corresponding IC50 (**B**, two-sided permutation t-test), as measured by Alamar blue. **E.** Cell growth of wild type and CTR1 KO cells in complete media and serum depletion after 48h as measured by total protein, normalized to the seeding cell number at time zero. Insert shows the percent difference in cell counts after the initial 48h. n=4 Average ± SEM. **F.** Normalized cell counts in wild type and CTR1 KO cells either in the absence or presence of 240 nM emetine for the indicated times. CTR1 clone KO20 was used for all experiments in **C-F**.

**Figure 6-1.**
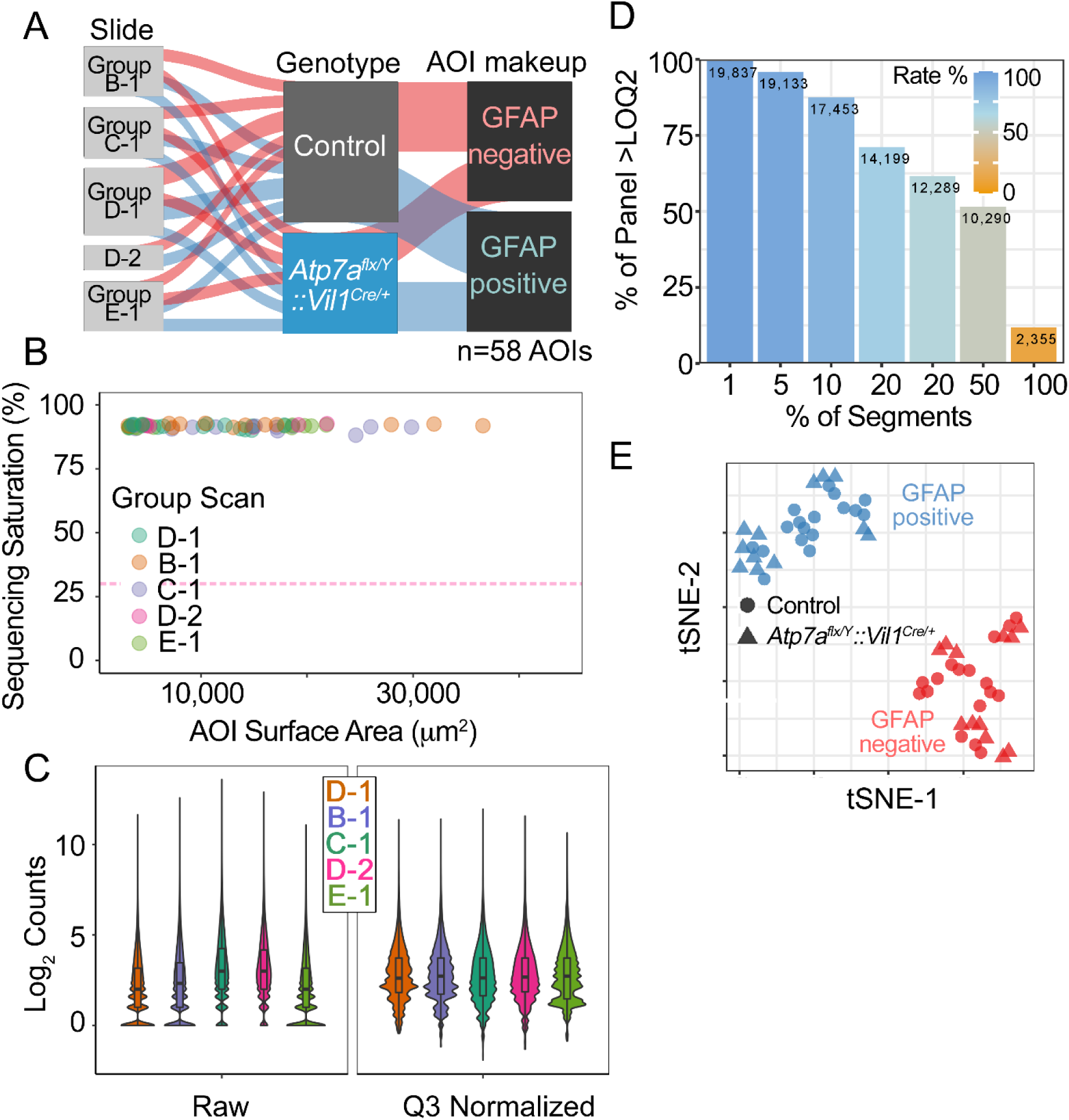
Spatial Transcriptomics Quality Controls and Descriptors. **A.** Sankey diagram depicts an overview of samples (n=58) showing their annotations. **B**. Sequencing quality as measured by saturation to ensure sensitivity of low expressor genes. Sequencing saturation measures sequencing depth for a sample defined as (1 – % Unique reads). Threshold above 50% is considered a reliable determination. 0 samples below the 50% threshold. **C**. Sequencing reads raw and normalized to the third quartile (Q3) to account for differences in cellularity, area of interest (AOI) size, or other variables. **D.** number of genes in different percentages of tissues expressed above LOQ2 (Limit of Quantitation), defined as the Geometric Mean of the negative probes, multiplied by the geometric standard deviation to the second power (^2). We assayed 19,963 genes across 58 AOIs. Of these 17,453 genes were expressed in 10% of AOIs and 10,290 genes expressed were expressed in 50% of AOIs. **E**. t-Distributed Stochastic Neighbor Embedding (t-SNE) of gene expression values per annotation. See Extended Data File 2 for raw data.

**Figure 6-2.**
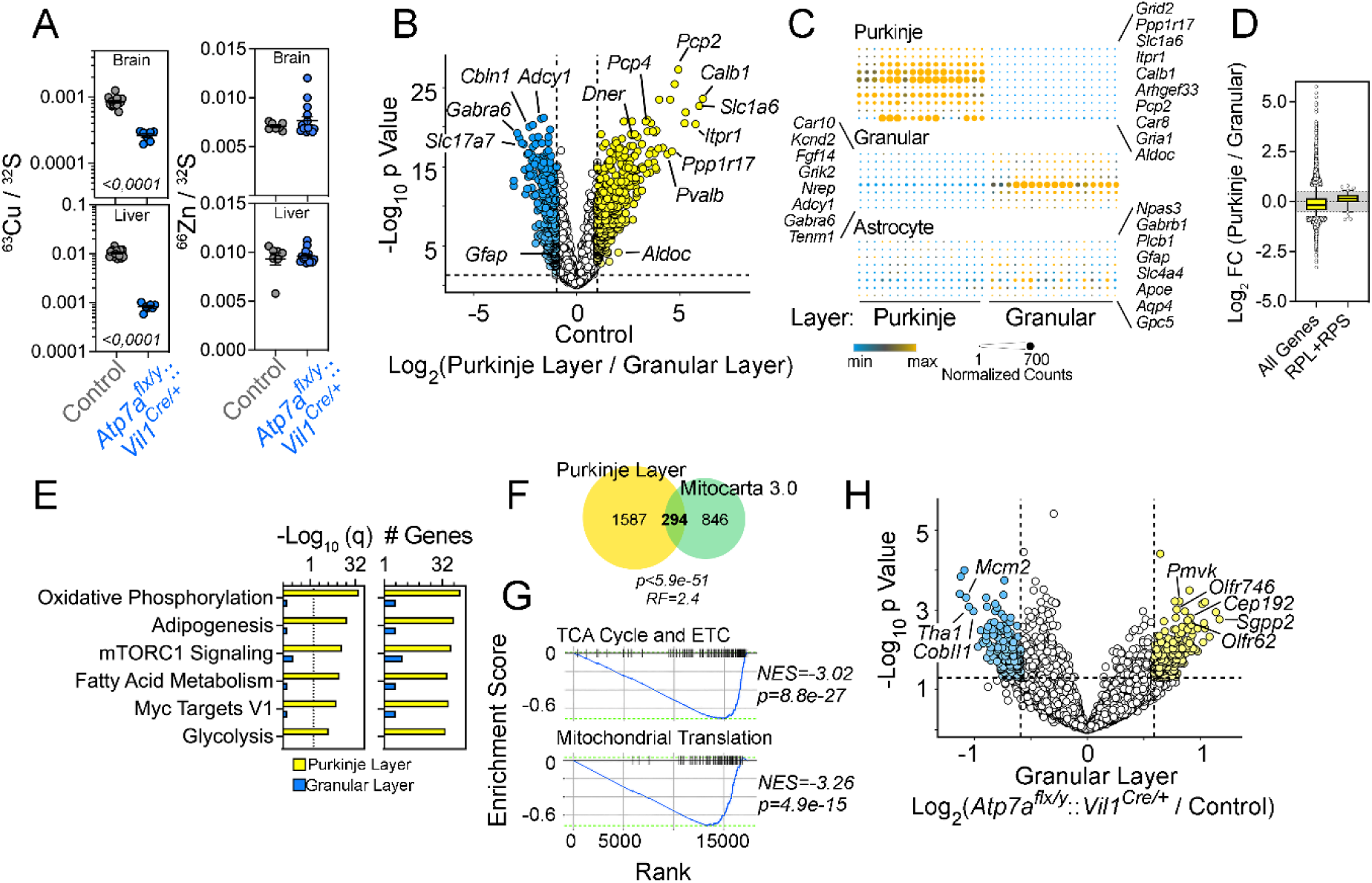
Cell type-specific gene expression in the cerebellar cortex in a presymptomatic Menkes mouse model. **A.** 63Cu and 64Zn quantification in the brains of *Atp7a^flx/Y^ :: Vil1^Cre/+^* mice at P10, normalized to 32S and analyzed by two-sided permutation t-test (italicized numbers represent p values). **B.** Volcano plot of mRNAs differentially expressed between the Purkinje and granular layer of control mice. **C.** Expression of selected cell type-specific markers (Kozareva et al., 2021). **D.** Differential expression of all transcripts compared to cytoplasmic ribosome subunits mRNAs. Whiskers correspond to 5-95 percentile, box represents 25 and 75 percentile, horizontal line marks the mean. Shaded area demotes ±1.5 fold difference. **E.** Gene ontology analysis of transcripts more highly expressed in Purkinje cells than granular layer cells. MSigDB was queried with the ENRICHR engine. Fisher exact test followed by Benjamini-Hochberg correction. **F.** 294 of the 1881 transcripts more highly expressed in wild-type Purkinje cells are annotated to the MitoCarta3.0 knowledgebase. **G.** GSEA and NES enrichment score of genes differentially expressed in wild-type Purkinje cells and granular layer cells reveal enrichment in metabolic ontologies. **H.** Volcano plot of mRNAs differentially expressed in the granular layer in mutant *Atp7a^flx/Y^ :: Vil1^Cre/+^* mice vs control. Yellow symbols mark genes with increased expression in the granular layer in mutant *Atp7a^flx/Y^ :: Vil1^Cre/+^* mice. See Extended Data File 2 for raw data.

**Figure 6-3.**
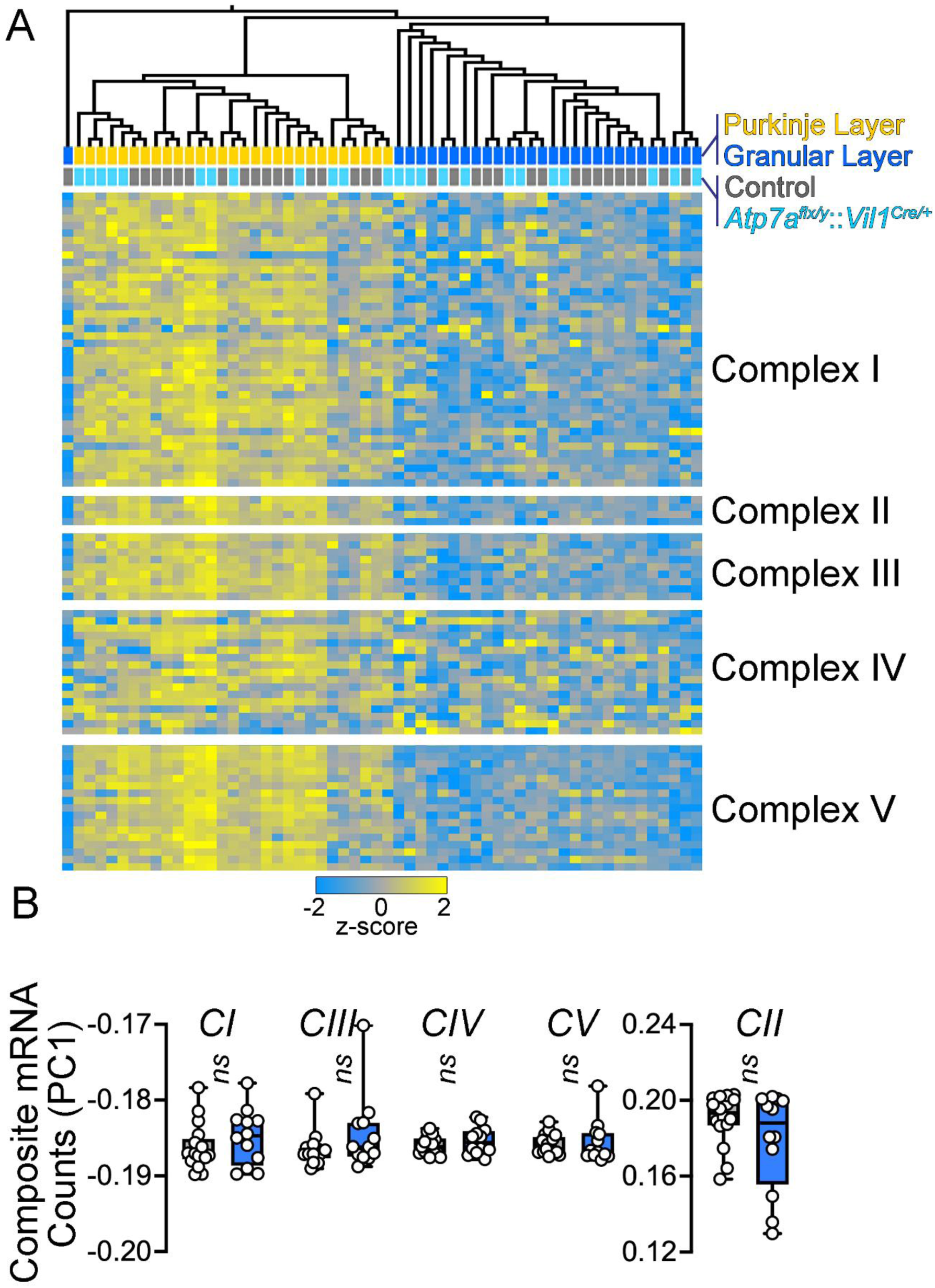
Respiratory complex subunit gene expression in a presymptomatic Menkes mouse model. **A.** Hierarchical clustering of all transcripts annotated to electron transport chain subunits according to MitoCarta 3.0 across genotypes and AOIs quantified. **B.** Principal component 1 of data presented in A for the Purkinje layer. Permutation t test. See Extended Data File 2 for raw data.

**Figure 7-1.**
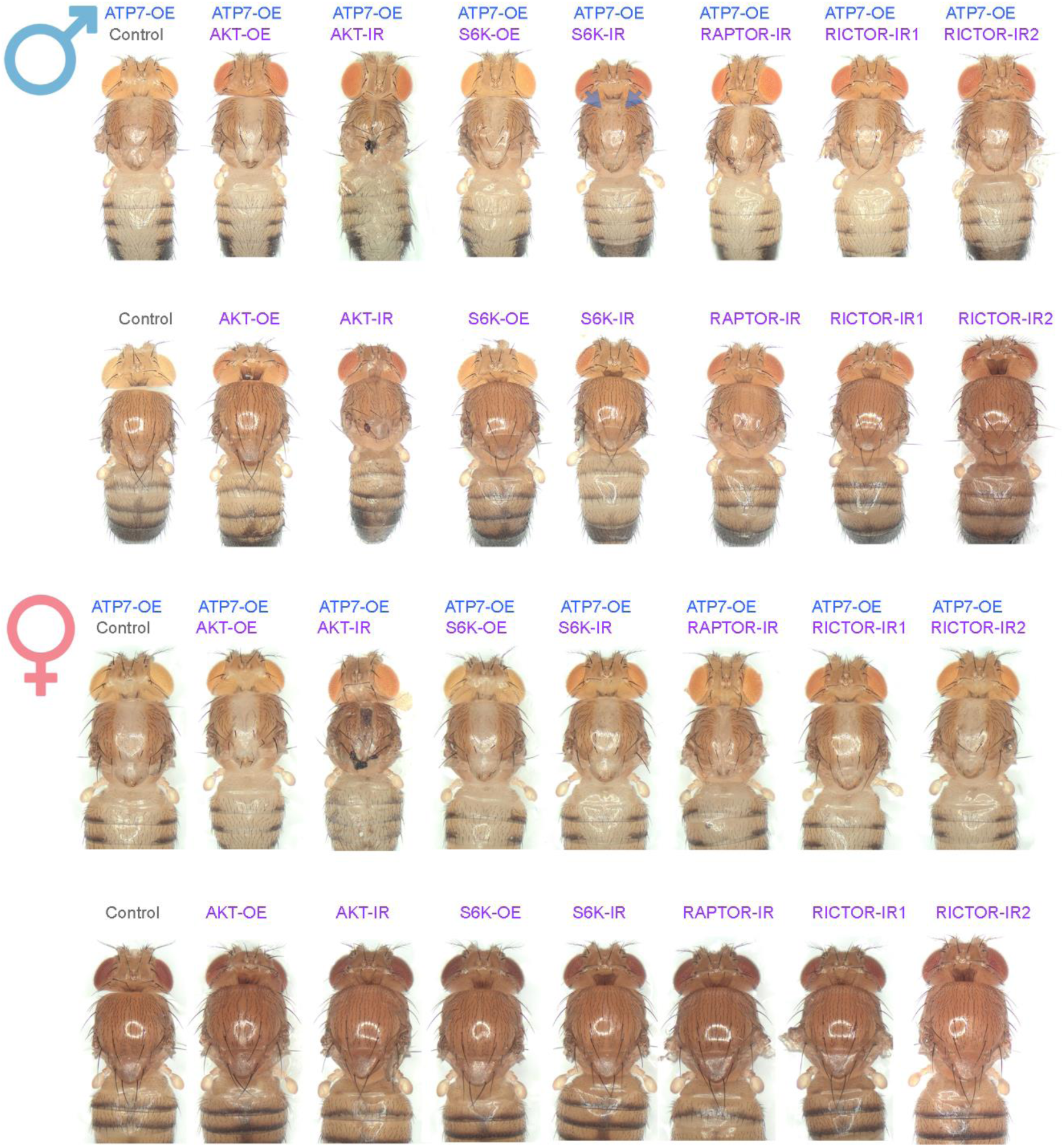
mTOR-Raptor-S6K pathway loss-of-function enhances copper-depletion phenotypes in *Drosophila* epidermis. Expression of wild type ATP7 and/or wild type or RNAi of different components of the mTOR pathway in the epidermis of the thoracic segment using the pnr-GAL4 driver in males and females (see Table 3 and Methods). Arrowheads point to ‘dimples’ in the dorsal aspect of ATP7-OE; S6K-IR males.

